# Primo: integration of multiple GWAS and omics QTL summary statistics for elucidation of molecular mechanisms of trait-associated SNPs and detection of pleiotropy in complex traits

**DOI:** 10.1101/579581

**Authors:** Kevin J Gleason, Fan Yang, Brandon L Pierce, Xin He, Lin S Chen

## Abstract

To provide a comprehensive mechanistic interpretation of how known trait-associated SNPs affect complex traits, we propose a method – Primo – for integrative analysis of GWAS summary statistics with multiple sets of omics QTL summary statistics from different cellular conditions or studies. Primo examines SNPs’ association patterns to complex and omics traits. In gene regions harboring known susceptibility loci, Primo performs conditional association analysis to account for linkage disequilibrium. Primo allows for unknown study heterogeneity and sample correlations. We show two applications using Primo to examine the molecular mechanisms of known susceptibility loci and to detect and interpret pleiotropic effects.

## Background

In the post-genomic era, genome-wide association studies (GWAS) have identified tens of thousands of unique associations between single nucleotide polymorphisms (SNPs) and human complex traits [1, 2]. Most of the trait-associated SNPs have small effect sizes and many reside in non-coding regions [3, 4], obscuring their functional connections to complex traits. It is known that trait-associated SNPs are more likely to also be expression quantitative trait loci (eQTLs) [5], thus many of these SNPs likely affect complex traits through their effects on expression levels and/or other “omics” traits. Extensive evaluations of genetic effects on omics traits such as gene expression [6], protein abundance [7], DNA methylation [8], histone modification [9, 10], and RNA splicing [11] have revealed an abundance of quantitative trait loci (QTLs) for omics traits (omics QTLs) throughout the genome. These findings suggest that integrating data from omics and multi-omics QTL studies with GWAS would help to elucidate functional mechanisms that underlie trait/disease processes. Moreover, the integrative analysis of omics and multi-omics traits would also enhance confidence in detecting true omics-associations while reducing false positive findings by observing co-occurrence of associations in multiple different data types and borrowing information across multi-omics data sources. The increasing availability of summary statistics for complex traits and omics QTL studies in many conditions and cellular contexts [6, 12, 13, 14] provides a valuable resource to conduct integrative analyses in a variety of settings and presents an unprecedented opportunity to gain a system-level perspective of the regulatory cascade, which may highlight targets for disease prevention and/or treatment strategies.

To integrate GWAS and omics QTL summary statistics, several methods have been proposed to identify trait-associated loci that share a common causal variant with 1-2 sets of omics QTLs (often referred to as “colocalization”) [15, 16, 17, 18]. There are also methods that have been proposed to directly test the molecular mechanisms through which genetic variation affects traits by integrating GWAS and eQTL summary statistics [19, 20]. These methods have identified known and novel candidate genes underlying psychiatric disorders [16], diabetes traits [18], obesity-related traits [15, 17, 19], and others. By applying the integrative methods to multi-omics data, some QTL pairs such as eQTL and methylation (me)QTL pairs have also been identified with evidence of a shared causal mechanism [16, 21]. Integrating studies of multiple complex and omics traits could produce a more comprehensive picture of how cellular processes contribute to variation in complex traits.

Compared to integrating GWAS with single omics QTL statistics, studying multi-omics QTLs increases the chances of detecting the regulatory mechanisms underlying trait/disease-associated SNPs. The effect of any particular SNP may be strong for some omics traits and weak or absent for others. For example, protein (p)QTLs exist for genes lacking an apparent eQTL [22], suggesting post-transcription regulation [23]. And there could be multiple different omics QTLs in a gene region with different functions. As another example, SNPs affecting RNA splicing (splicing QTLs) may not be eQTLs in a gene region [11]. Moreover, QTL effects may vary across molecular phenotypes [24], tissue types [6], cell types [25, 26], or other contexts [27, 28]. For example, lead SNPs for eQTLs (eSNPs) often vary by tissue type [6]. Jointly analyzing the omics QTL association summary statistics to more than one type of omics trait from different conditions/studies could yield a more complete portrait of the regulatory landscape. Given the increasing availability of summary statistics for omics QTLs from different studies/conditions/cell-contexts, novel methods and tools are needed to integrate GWAS with many relevant sets of omics QTL summary statistics for an improved understanding of the mechanisms of trait-associated SNPs.

Jointly analyzing more than three complex and omics traits can also be viewed as an approach for identifying shared mechanisms that underlie multiple complex traits – pleiotropic effects. Pleiotropy is ubiquitous in the genome [29, 30]. Since pleiotropic effects often occur among related diseases and traits [31, 32, 33], shared mechanisms are likely to exist. By integrating omics QTL summary statistics from multiple trait-relevant tissue types with GWAS statistics, one can also boost power in detecting pleiotropic effects while simultaneously providing mechanistic interpretations.

It is desirable to develop new methods that can integrate multiple (i.e. more than three) sets of GWAS statistics and omics QTL statistics from different conditions/studies while accounting for study heterogeneity, potential sample correlations and linkage disequilibrium (LD). Additionally, as the number of traits/studies/conditions being considered grows, it will be more likely to detect joint associations by chance, necessitating proper multiple testing adjustment. To address those challenges, in this work we develop a method to integrate summary statistics from different studies to examine the genetic effects on multiple complex and omics traits, and implement the method in an R package – Primo (Package in R for Integrative Multi-Omics association analysis). Figure 1A provides an overview of the algorithm. Different than the traditional meta-analysis approaches that also take summary statistics as input, Primo is flexible in many aspects: it allows unknown and arbitrary study heterogeneity and can detect coordinated effects from multiple studies while not requiring the effect sizes to be the same; it allows the summary statistics to be calculated from studies with independent or overlapping samples with unknown sample correlations; and it is not an omnibus test for association, but rather can be used to calculate the probability of each SNP belonging to each type (or groups) of interpretable association patterns (e.g. the probability of a trait-associated SNP also being associated with at least one/two cis omics-traits). For gene regions harboring known susceptibility loci, the conditional association analysis of Primo examines the conditional associations of a known trait-associated SNP with other complex and omics traits adjusting for other lead SNPs in a gene region. It moves beyond joint association towards causation and colocalization, and provides a thorough inspection of the effects of multiple SNPs within a region to reduce spurious associations due to LD (Figure 1B). We conduct extensive simulations to evaluate the performance of Primo under various scenarios in analyzing multiple sets of summary statistics from studies with correlated samples. We apply Primo to examine the omics trait association patterns for known SNPs associated with breast cancer risk by integrating multi-omics QTL summary statistics from the Genotype-Tissue Expression (GTEx) project [6] and The Cancer Genome Atlas (TCGA) [34] with GWAS statistics from The Breast Cancer Association Consortium (BCAC) [35]. We also apply Primo to detect SNPs with pleiotropic effects to two complex traits in gene regions harboring susceptibility loci for at least one trait, while also providing mechanistic interpretations by integrating publicly available GWAS summary statistics [36, 37, 38, 39] with multi-tissue eQTL summary statistics from GTEx.

**Figure 1.**
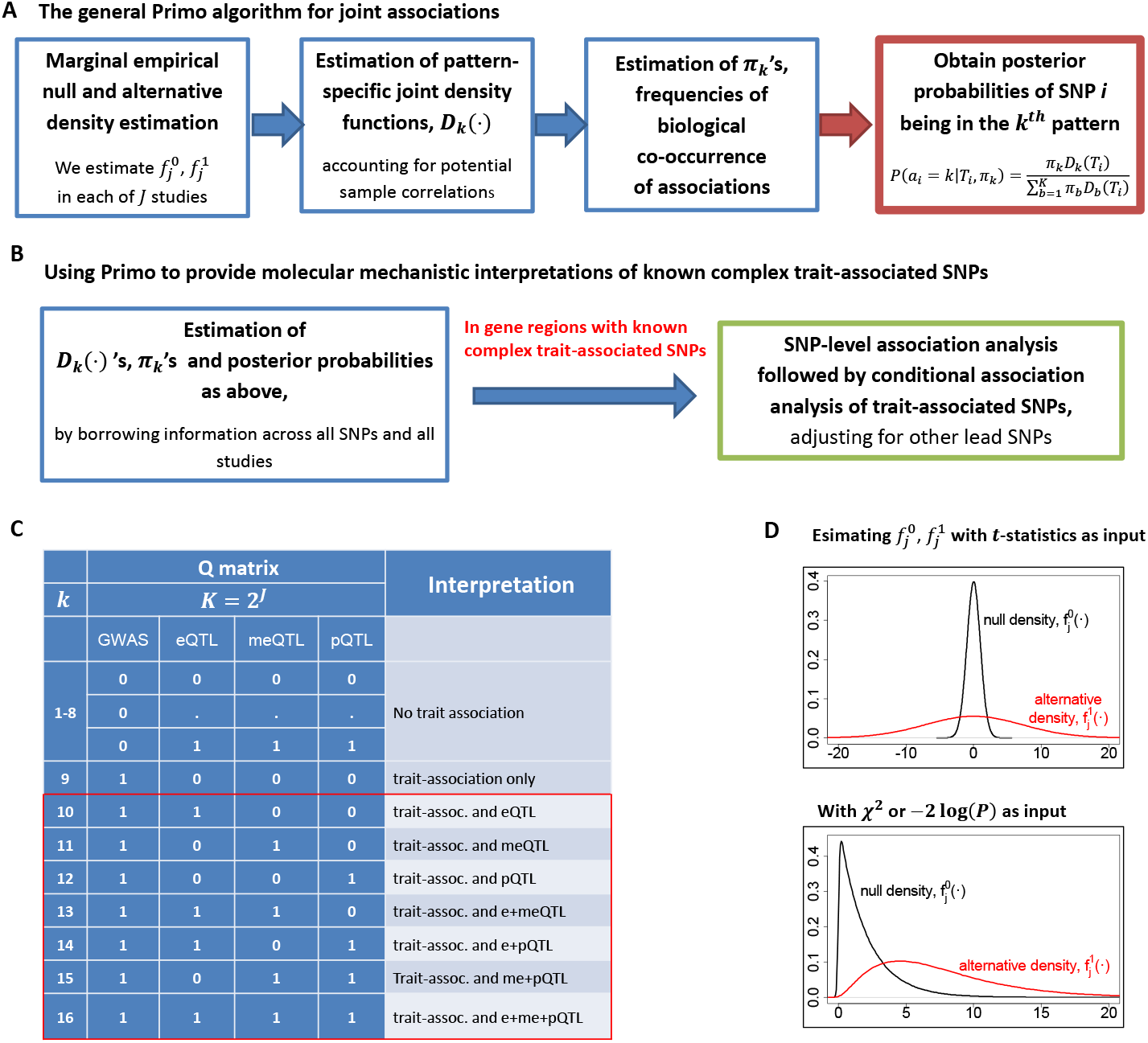
An overview of Primo and an illustration of interpretations of results. (A) The main steps of the Primo algorithm for assessing joint associations. (B) Steps of the Primo algorithm to provide mechanistic interpretations of known complex trait-associated SNPs (C) An example – the **Q** matrix and interpretations of association patterns for an analysis of a complex trait, eQTL, meQTL and pQTL studies for *j* = 1, 2, 3 and 4, respectively. The red box shows how association patterns can be collapsed into groups of interest (here, summing probabilities across the patterns in the red box would yield the probability of association with the complex trait and at least one omics trait). (D) An example of the estimated marginal null and alternative densities of a moderated *t*-distribution (top) and *χ*^2^ or −2log(*P*) values (bottom) for a study *j*.

## Results

### Primo as a general framework for assessing joint associations across data types

Here we first introduce the general Primo association framework (Figure 1A) and then discuss the tailored development in using Primo to provide mechanistic interpretations of known trait-associated SNPs (Figure 1B), moving from association to colocalization. As a general integrative association method, Primo takes as input multiple sets of association summary statistics from different studies of different data types. The multiple sets of summary statistics could be one set of GWAS statistics and multiple sets of omics/multi-omics QTL statistics, or two or more sets of GWAS statistics of related traits and multiple sets of omics/multi-omics QTL statistics from trait-relevant tissue types, or could even be from studies beyond the complex and omics trait-associations of germline variation.

Consider an *m* × *J* matrix of association statistics, **T**, consisting of the summary statistics for the associations of *m* SNPs with *J* types of traits from *J* studies with independent or correlated samples. Note that here a “study” refers to a study of SNPs’ associations to a particular trait in a particular condition/cell-type/tissue-type. For each SNP (here a row in the matrix **T**), the underlying association status to the *j*-th (*j* = 1,…, *J*) trait is binary. Considering all SNPs in the genome, there are a total of *K* = 2*^J^* possible association patterns to *J* traits. We use a *K × J* binary matrix, **Q**, to denote all of the possible association patterns. And *q_kj_* = 1 implies *the presence of association* with the *j*-th trait in the *k*-th association pattern, and *q_kj_* = 0 implies *no association*.

For each SNP *i*, there must be one and only one true underlying association pattern. Primo calculates the probability of a given SNP being in each of the *K* mutually exclusive association patterns by borrowing information across SNPs in the genome and across *J* traits. More specifically, let *a_i_* denote the true association pattern for SNP *i*. Then the probability that SNP *i* belongs to association pattern *k* is given by:

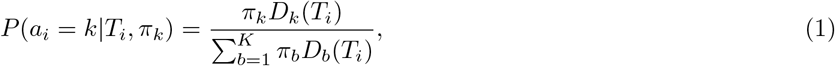

where *T_i_* is a vector of *J* association statistics and is also the *i*-th row in the **T** matrix, *π_k_* represents the overall proportion of SNPs in the genome belonging to the *k*-th association pattern (*k* = 1,…, *K*), and *D_k_*(·) is the multivariate density function of *J* sets of statistics, conditioning on the *k*-th association pattern. Here *π_k_* captures the biological co-occurrence frequency of the *k*-th association pattern in the genome, with 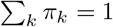. For example, in Figure 1C, *π*_16_ is the proportion of SNPs in the genome that are associated with all of the three omics traits and the complex trait.

In estimating a mixture distribution of *K* components, the performance of estimation and subsequent inference depend on how well different mixing components separate from each other. When *K* is moderate to large, it is challenging to simultaneously estimate the distributions of mixing components (*D_k_*’s) and the mixing proportions (*π_k_* ‘s). Different from previous work [40], Primo first estimates the pattern-specific multivariate density function *D_k_* for each of the association pattern by borrowing information across SNPs and traits. See Methods section for detailed estimation procedures when *J* sets of association statistics were calculated from independent or correlated samples. Then Primo estimates *π_k_*’s via the Expectation-Maximization algorithm [41]. When *D_k_*’s are reasonably-estimated, the one-step estimates of *π_k_*’s can well capture the overall proportions of different association patterns and there is no need to re-iterate and re-estimate *D_k_*’s and *π_k_*’s. Based on (1), we can obtain the posterior probabilities of SNP *i* being in each of the *K* possible association patterns.

### Mechanistic interpretations of trait-associated SNPs via Primo conditional association analysis in gene regions harboring susceptibility loci

In order to elucidate the molecular mechanisms of known trait-associated SNPs, one may examine the omics trait associations of those SNPs by integrating GWAS and omics QTL summary statistics. However, a major challenge in such analyses is the complex LD structure among SNPs in the same gene region.

To assess whether a GWAS SNP is associated with omics traits not due to it being in LD with other lead omics SNPs, we propose to conduct conditional association analysis within gene regions harboring susceptibility loci, with summary statistics of the GWAS SNP and other lead omics SNPs as input. Here we consider a GWAS SNP *i* of interest and a set of lead omics SNPs *I* in the gene region, where *I* is a set of indices. We can model the joint association statistics for SNPs *i* and *I* in study *j* using a multivariate normal distribution, and further calculate the conditional density functions of SNP *i* adjusting for other lead omics SNPs given their most plausible association patterns. See Methods for details. Then with the estimated *π_k_*’s, we can assess the probabilities of associations for SNP *i* in (1). Figure 2 shows a conceptual illustration of the conditional association analysis. If the GWAS SNP is an independent meQTL and pQTL, it remains associated with methylation and protein after adjusting for other lead SNPs in the region; and if the GWAS SNP is associated with cis-expression levels because it is in LD with the lead eSNP, it will be no longer significantly associated with expression after adjusting for the lead eSNP. With conditional association analysis, we can reduce spurious associations due to LD.

**Figure 2.**
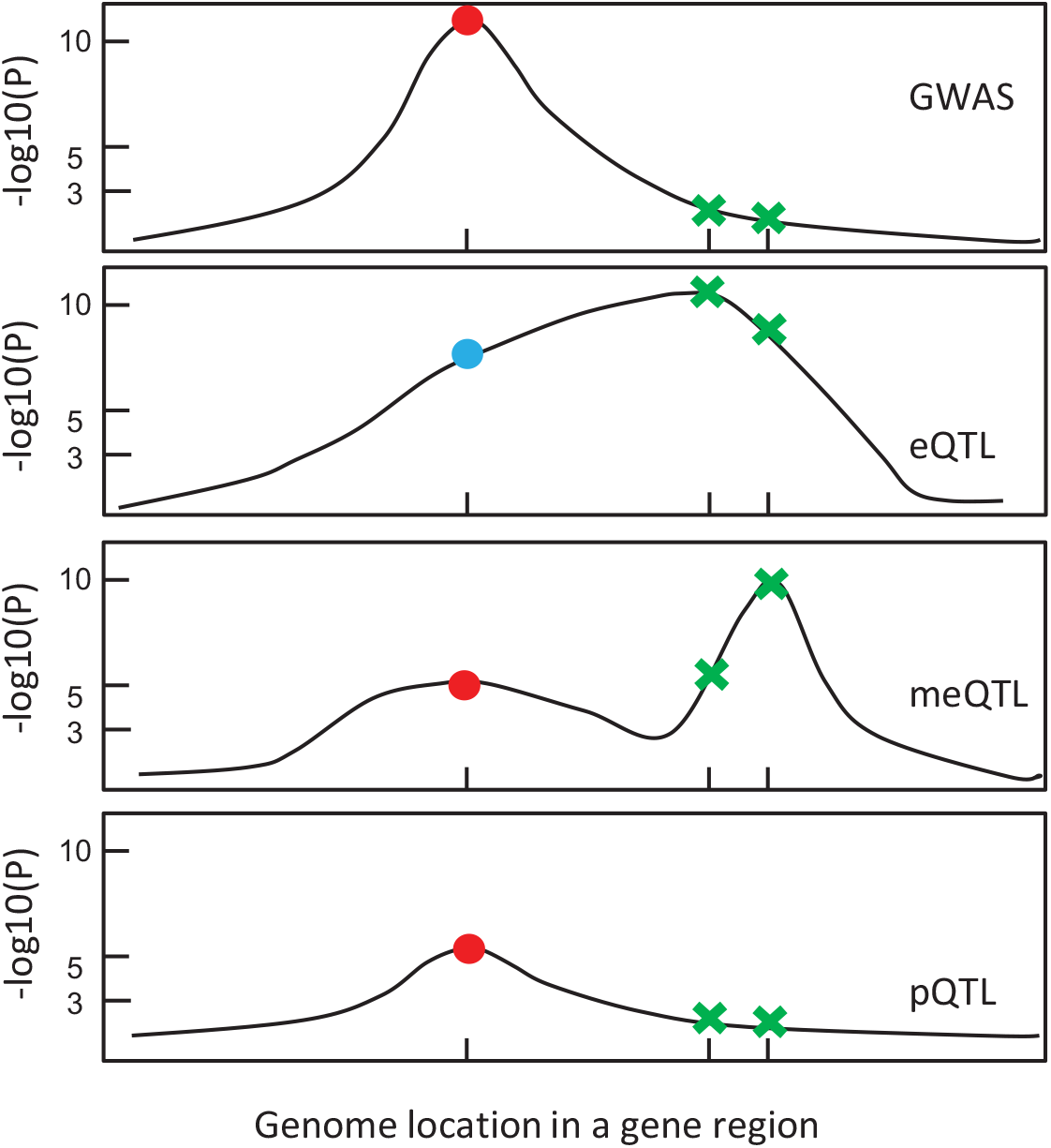
A conceptual illustration of the conditional association analysis of Primo. Consider a joint analysis of GWAS summary statistics and summary statistics of eQTL, meQTL and pQTL. In a gene region harboring trait-associated SNPs, there is a GWAS SNP of interest (red/blue dot) and two other confounding SNPs – the lead SNPs for eQTL and meQTL (green cross). Before conditional association analysis, the GWAS SNP is estimated to be associated with cis expression, methylation and protein levels. After adjusting for the two lead omics SNPs, the GWAS SNP is no longer associated with cis expression levels (blue dot) but is still estimated to be a me- and pQTL.

An advantage of Primo is that one may collapse many association patterns based on biological interpretations and obtain the posterior probabilities of groups of patterns of interest by summing over the probabilities of those mutually exclusive patterns. As illustrated in Figure 1C, when *J* = 4, there are 16 possible association patterns. We may collapse the association patterns into interpretable groups. For example, here we are interested in the trait-associated SNPs that are also associated with at least 1 omics trait. And we can obtain the probability estimate by summing over the posterior probabilities of patterns 10-16. For a (collapsed) pattern of interest, we can also calculate the estimated false discovery rate (FDR) [42] for multiple testing adjustment:

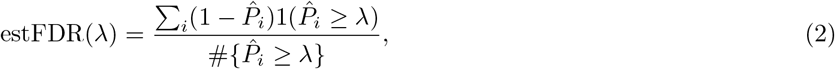

where λ is the probability threshold and 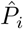 is the estimated probability of SNP *i* being in the (collapsed) pattern of interest.

As a summary, to elucidate the molecular mechanisms of trait-associated SNPs, we first obtain the estimates of key parameters (*π_k_*’s, *D_k_*’s) by borrowing information across SNPs and across traits/studies. Then in each gene region harboring known trait-associated SNPs, we conduct a SNP-level association analysis to all traits for all SNPs in the gene region, followed by a conditional association analysis for each GWAS SNP accounting for LD with other lead omics SNPs. If a GWAS SNP is no longer associated with a particular omics trait after conditioning on the lead omics SNPs, we will not consider it as a causal SNP for that omics trait. Estimated FDR can be calculated as described.

In the Methods section, we also discuss extensions of Primo, with *P*-values as input or when the number of traits being considered is large (> 15). We implemented Primo in Rcpp. It is computationally efficient and can analyze the associations of 30 million SNPs to five sets of complex and omics traits within 20 minutes on a single machine with 32 GB of memory and a 3 GHz processor.

### Simulation studies to evaluate the performance of Primo

We evaluated the performance of Primo in a variety of simulated scenarios. In each scenario, we simulated the test statistics for associations of SNPs with *J* traits. Test statistics under the null hypothesis of no association were simulated from a standard normal distribution; test statistics under the alternative were simulated from a normal distribution with mean 0 and standard deviation of 10 (allowing effect sizes to be positive or negative). The simulated data structure and test statistic distributions mimic what we have observed in the eQTL data from GTEx. For each simulated dataset, we ran two versions of the Primo algorithm using *t*-statistics and *P*-values as input, denoted as Primo (*t*) and Primo (*P*), respectively. We repeated each simulation 100 times and compared the performance of the two versions of Primo versus competing methods (if applicable).

#### Accurate estimation of proportions (π) even for very sparse joint associations

It is known to be challenging to estimate *π_k_*’s when associations are sparse, i.e., *π_k_*’s being very close to zero for patterns with associations. In Scenarios 1a and 1b, we showed that in analyzing independent and correlated sets of summary statistics, respectively, Primo can well estimate the *π_k_*’s despite very sparse associations. In each scenario, we simulated test statistics for *J* = 3 traits for 10 million SNPs, first under independence and then with pairwise (Pearson) correlation of 0.3 between each set of statistics. In Scenario 1a, we simulated true *π_k_* = (7 × 10^−4^, 2 × 10^−4^, 1 × 10^−4^) for SNPs being associated with only one, exactly two, and all three traits, respectively. Scenario 1b simulated even sparser associations for the third trait, with π_k_ = (7 × 10^−6^, 2 × 10^−6^, 1 × 10^−6^) for SNPs being associated with only the third, the third and first or second, and all three traits, respectively. Table 1 shows true *π_k_*’s, and the average estimates for *π_k_*’s by Primo based on *t*-statistics or *P*-values. In *Supplemental Materials*, we also show the performance of estimation of *π_k_*’s when the marginal alternative proportions 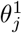’s are mis-specified. As shown, Primo estimates the *π_k_*’s with reasonable accuracy even when the associations are very sparse and when the marginal alternative proportions 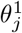’s are under-specified.

**Table 1.**
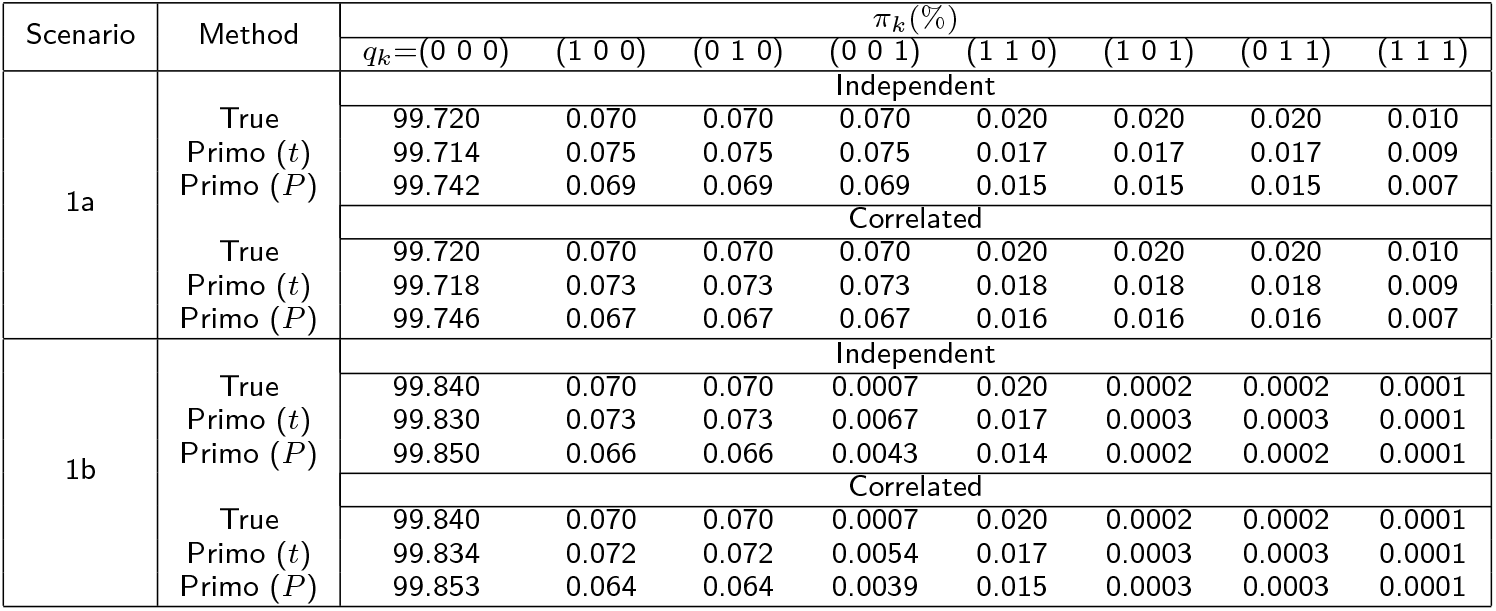
Average estimates of 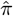. Scenario 1a simulates sparse associations for *J* =3 traits. Scenario 1b simulates even sparser associations for the third trait.

#### Comparison with existing methods for jointly analyzing associations to three traits

In Scenario 2, we simulated correlated test statistics with pairwise correlations of 0.3 among *J* = 3 traits for 1 million SNPs. *π_k_* = 1 × 10^−3^, 5 × 10^−4^, 5 × 10^−4^ for the patterns where SNPs are associated with only one, exactly two, and all of the three traits, respectively. Here we compared the true and estimated FDRs and power to detect associations to all three traits and to at least one trait, based on Primo versus two competing methods, “moloc” [16] and Fisher’s method [43]. The results with correctly specified, under-specified and over-specified marginal non-null proportions 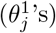 are shown in Table 2. When 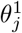’s are well-specified (Scenario 2a in Table 2), Primo nicely controlled the FDR even in the presence of unknown study/sample correlations – highlighting one advantage of Primo in integrating potentially correlated multi-omics data. Note that moloc and Primo are not directly comparable as moloc aims to detect the true causal variant in a gene region while Primo first identifies SNPs’ joint associations to multiple traits and then reduces spurious associations due to LD. Nevertheless, we grouped sets of 100 SNPs together to form “regions” to allow for some comparisons between Primo and moloc. For half of the gene regions, there are two independent multi-omics QTLs; and for the other half, there is only one multi-omics QTL in a region. Since moloc does not output the posterior probabilities for all SNPs in every association pattern, we are only able to compare the power and FDR of Primo versus moloc in detecting associations to all three traits. We observed that Primo generally enjoys substantial power improvement, which is not surprising because the goal of moloc is more restrictive. As shown in Table 2, the estimated FDR (estFDR) is very close to the true FDR for Primo. Fisher’s method, as a combination method for testing omnibus hypotheses, can only be used to detect SNPs with associations to at least one trait, and is not applicable to detect associations to all traits. The true FDR [42, 44] for Fisher’s method calculated from the nominal *P*-values are not well controlled due to correlations among test statistics, as expected. At similar power levels, the true FDRs of Fisher’s method are also much higher than those of Primo.

**Table 2.**
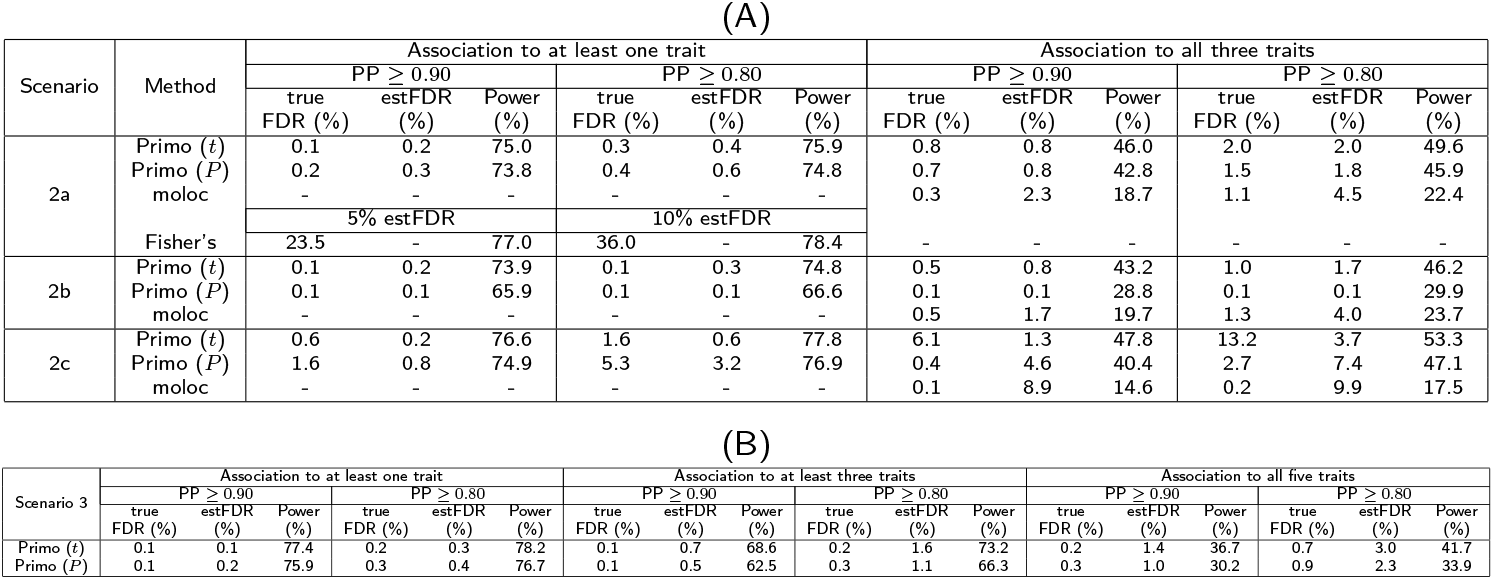
Simulation results evaluating the performance of Primo. PP := posterior probability; estFDR := estimated FDR.(A) When *J* = 3 with correlated samples, we compared Primo versus moloc and Fisher’s method in detecting associations to at least 1 trait and associations to all traits and when parameters are correctly, under- and over-specified. (B) When *J* = 5 with correlated samples, we evaluated the performance of Primo.

In this simulation, the true 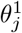’s are 2.5 × 10^−3^. In Scenario 2b, we under-specified 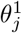 to be 2.5 × 10^−4^. As shown in Table 2(A), although power might decrease to some extent, the FDRs are reasonably controlled. In Scenario 2c, when 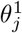’s are overspecified as 2.5 × 10^−2^, we observed slightly inflated FDRs. As such, we suggest to obtain reasonable estimates for 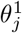’s based on the current data and the literature, or under-specify 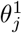’s to be more conservative.

#### The performance of Primo in jointly analyzing more than three traits

In Scenario 3, we simulated correlated test statistics for associations to five traits for 1 million SNPs with pairwise study-study correlations of 0.3. *π_k_* = 5 × 10^−4^, 2 × 10^−4^, 1 × 10^−4^ for the patterns where SNPs are associated with one to two, three to four, and all of the five traits, respectively. Results are presented in Table 2(B). Overall, Primo yields good control of FDRs and high power in detecting various patterns of joint associations, even for a moderately large number of sets of summary statistics and in the presence of study correlations.

#### Evaluation of the performance of Primo conditional association analysis accounting for LD and sample correlations

In this section, we simulated association statistics for correlated SNPs in moderate to high LD and evaluated the performance of the proposed conditional association approach in the presence of both LD and sample correlations. We simulated the matrix of association statistics **T** for 1 million SNPs with *J* = 4 traits as the sum of two statistics matrices, **T**^(1)^ and **T**^(2)^, where **T**^(1)^ was simulated to impose LD structures and **T**^(2)^ was simulated to impose sample correlations. To simulate **T**^(1)^, we created 10^5^ LD blocks with 10 SNPs in each. The statistics for SNPs from different LD blocks are independent, whereas statistics for SNP *i* and SNP *j* from the same block have a correlation 0.95^|*i−j*|^ in the same study. Among 5% of the LD blocks, the true underlying association patterns are (1, 0, 1, 0), (1, 1, 0, 0) and (0, 1, 1, 1) for SNPs 4, 5, and 6, respectively; and (0, 0, 0, 0) for the rest. In those blocks, the marginal null distribution is standard normal and the marginal alternative distribution is N(3.5, 1). Those LD blocks represent gene regions with no SNP truly associated with all traits but with multiple SNPs in LD with different association patterns. We further simulated another 5% of the LD blocks, where the true underlying association patterns are (0, 0, 0, 1), (0, 1, 0, 0) and (1, 1, 1, 1) for SNPs 4, 5, and 6, respectively; and (0, 0, 0, 0) for the rest. In those blocks, the marginal alternative distributions for SNPs 4 and 5 are N(2, 1), and are N(3.5, 1) for SNP 6. Those LD blocks represent gene regions with one true causal SNP associated with all traits as well as two confounding SNPs in high LD with it. For the remaining 90% of the LD blocks, all the statistics are generated under the null. In **T**^(2)^, correlated statistics were generated with all pairwise correlations of 0.6 among traits and were all simulated under the null with standard deviations being 0.2. The resulting matrix **T** = **T**^(1)^ + **T**^(2)^ has both correlated rows and correlated columns.

We applied Primo with **T** as input to identify SNPs associated with all traits. For each SNP detected as significant at the probability cutoffs of 0.8 and 0.9, we further conducted conditional association analysis. In each LD block, SNPs 4, 5 and 6 served as the lead/confounding SNPs for each other. For instance, in testing the associations for SNP 4, we consider SNPs 5 and 6 as the lead SNPs in the region and adjusted them. For the rest of the SNPs in the block, SNPs 5 and 6 were used as the lead SNPs and were adjusted in the conditional analysis. The SNPs that no longer have the highest probabilities in the pattern of (1, 1, 1, 1) after conditional association analysis were not considered to be positive findings. In the calculations of the FDR’s, we use the same denominators before and after conditional association analysis for fair comparison. That is, the denominators are the number of identified SNPs with associations to all traits at a given cutoff before the conditional associating analysis. After conditional analysis, the numerator (i.e. # false positive) of the true FDR is the number of SNPs that are not truly associated to all traits, yet continue to show the highest probability in the pattern of (1, 1, 1, 1) after conditional association analysis. In the calculation of the numerator of the estimated FDR, for each SNP *i* that is no longer significant after conditional analysis, its contribution to the numerator 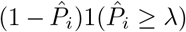 in the formula (2) is corrected to be 1 since we considered it as an estimated false discovery.

As shown in Table 3, when SNPs are in LD, we observed some slightly inflated FDRs without conditional association analysis even when 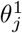’s are correctly specified (Scenario 4a). In contrast, after accounting for LD, true FDRs are reduced and are well controlled by the estimated FDRs. In Scenario 4b and 4c, we under-specified and over-specified 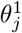’s. Overall, Primo after conditional association analysis could yield nice control of FDR and maintain good power in all scenarios.

**Table 3.**
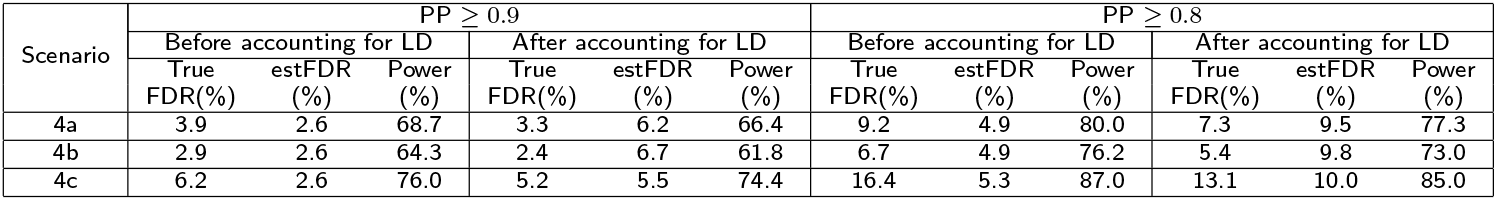
Comparison of results before and after conditional association analysis. PP := posterior probability

### Application I: Understanding the mechanisms of breast cancer susceptibility loci

With over 100,000 breast cancer cases and a similar number of controls from a total of 78 breast cancer studies, BCAC [35] has recently reported 174 common genetic variants associated with breast cancer risk. In order to understand the underlying mechanisms of those susceptibility risk loci and their potential cis target genes, a recent study [45] conducted cis-eQTL analysis using both normal and tumor breast transcriptome data and identified multiple genes likely to play important roles in breast tumorgenesis.

In addition to transcription, SNPs may affect cis-epigenetic features, protein abundances, and other omics traits. Functional relationships may exist among those omics traits. Therefore, we propose to jointly examine the susceptibility risk loci and their effects on multiple omics traits in tumor and normal tissues in order to better understand the mechanisms through which risk-associated SNPs acts in different conditions. Moreover, this analysis will enhance our understanding of the regulatory cascade and their roles in breast tumorigenesis. The regulatory SNPs with “cascading effects” [22, 46] on gene regulation and downstream gene products are of particular interest.

In this work, we applied Primo to integrate GWAS summary statistics from BCAC for all SNPs in the genome with the eQTL, meQTL, and pQTL association summary statistics obtained from 1012, 702, and 74 breast tumor samples, respectively, from TCGA [34, 47] (see *Supplemental Materials*) and eQTL summary statistics obtained from 397 normal breast mammary samples from GTEx [6]. A total of 161 of the GWAS SNPs reported by Michailidou, *et al.* (2017) reached genome-wide significance (*P* < 5 × 10^−8^) in the meta-analysis. And there are 157 of these SNPs with MAF > 1% in TCGA data. Note that one SNP could be mapped to multiple genes and multiple CpG sites. We assessed the probabilities of 32 (2^5^, for GWAS and 4 omics QTLs) association patterns for each SNP-gene-CpG-protein quartet. In the conditional association analysis of gene regions harboring at least one GWAS SNP, we selected the lead SNP for each omics trait in the region and adjusted for any lead SNP outside a 5kb distance of and with LD *R*^2^ < 0.9 with the GWAS-reported SNP (those with *R*^2^ > 0.9 or within 5kb were considered likely to share a causal variant or too close to assess individual associations, respectively).

At the 80% probability cutoff and after conditional association analysis (estimated FDR of 3.8, 6.2, 12.2, and 8.5%), there were 49, 18, 7 and 1 susceptibility loci associated with at least 1, 2, 3 or 4 omics traits, respectively. The three GWAS SNPs (rs11552449, rs3747479, and rs73134739) in the three genes (*DCLRE1B, MRPS30*, and *ATG10*, respectively) reported in Guo, *et al.* (2018) [45] had high probabilities of being an eQTL in both tumor and normal tissues (with probabilities of 59.7, >99.9, and >99.9%, respectively). In the *KLHDC7A* gene region, the GWAS SNP rs2992756 (indicated by red dot in Figure 3) is associated with the expression, methylation and global protein abundance levels of the cis-gene *KLHDC7A*. Figure 3 shows the plot of −log_10_ *P*-values of associations to breast cancer risk and the three omics traits (with expression traits in both tumor and normal tissue types) of *KLHDC7A* for the SNPs in the gene region. Note that the GWAS SNP is only moderately associated with the gene expression levels in the normal GTEx breast tissue with a *P*-value of 0.0034, highlighting the need to study omics QTLs under different conditions.

**Figure 3.**
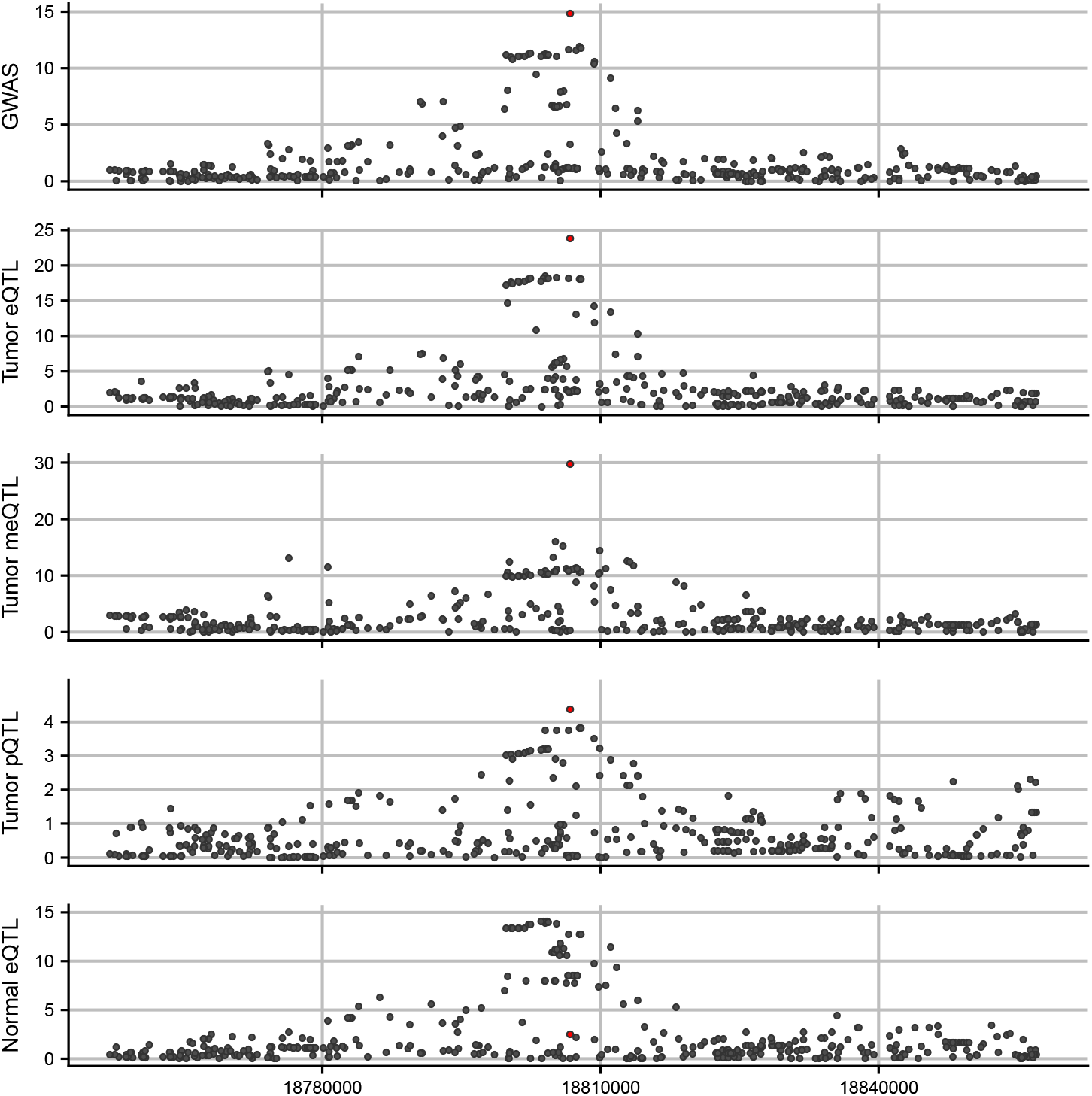
An example of a known breast cancer susceptibility locus being associated with multi-omics traits. Primo estimated a high probability (94.2%) of SNP rs2992756 being associated with all four omics traits. Here shows the −log_10_ (*P*)-values by position on Chromosome 1 in the region of the gene *KLHDC7A* for all SNPs including the breast cancer susceptibility locus (rs2992756, red dot) in GWAS (top panel) and eQTL, meQTL and pQTL analyses in tumor tissue (the next three panels, respectively) and eQTL analysis in normal tissue (bottom panel) for the gene and protein KLHDC7A and CpG site cg05040210.

Due to limited sample sizes (74) in the pQTL analysis, only 1 out of the 157 examined breast cancer susceptibility loci was associated with cis-protein abundance levels with high confidence, although the cis-gene expression levels and cis-protein abundances for those loci were often highly correlated with an averaged (Pearson) correlation coefficient of *r*=0.398 and a median of *r*=0.414. There were 21 out of 157 susceptibility loci uniquely associated with cis-methylation levels but not expression levels in either tumor or normal tissue, echoing a recent work showing both unique and shared causal mechanisms of epigenome variations and transcription [21]. This also shows that the integration of GWAS and multi-omics traits can provide additional insights in understanding the complex and dynamic mechanisms.

### Application II: Detecting SNPs with pleiotropic effects and elucidating their mechanisms

Many genetic variants are associated with more than one complex trait [48, 29, 30]. Identifying such pleiotropic variants and elucidating the molecular mechanisms which underlie these multi-trait associations may enhance our understanding of the etiology of complex traits and provide additional insights into clinical treatment development [48]. In this section, we applied Primo to detect SNPs with pleiotropic effects to two complex traits in gene regions harboring susceptibility loci for at least one trait, and provide mechanistic interpretations by integrating pairs of publicly available complex-trait GWAS summary statistics with eQTL association summary statistics obtained from trait-relevant tissue types in the GTEx project.

We applied Primo to height [37] and body mass index (BMI) [38] GWAS summary statistics from the GIANT consortium (sample size > 250, 000) with eQTL summary statistics in subcutaneous adipose (*n* = 581) and skeletal muscle (*n* = 706) tissues from GTEx for all SNPs in the genome. There are 697 height-associated SNPs reported by Wood, *et al.* [37], and 623 of them have reached genome-wide significance threshold (5 × 10^−8^) while the others are significant in the conditional analysis. There are 97 BMI-associated SNPs reported by Locke, *et al.* (2015) [38], 77 of which reached the genome-wide threshold in the sex-combined analysis. Out of the 700 genome-wide significant SNPs for either trait, 683 were present in both sets of GWAS summary statistics and could be mapped to GTEx SNPs in cis with at least one gene measured in both tissue types. At the 80% probability cutoff and after conditional association analysis accounting for LD, 32 SNPs were associated with both complex traits (estimated FDR of 12.4%). Of these, 18 were associated with expression of at least one gene in at least 1 tissue (estimated FDR of 13.6%) and 12 were associated with expression of at least one gene in both tissues (estimated FDR of 12.5%). Furthermore, 13 of the SNPs were associated with the expression of multiple genes, highlighting the possibility that pleiotropic SNPs may affect multiple complex traits through their co-regulation of multiple genes.

To validate the 32 pleiotropic SNPs being associated with both height and BMI, we used GWAS summary statistics from the UK Biobank [39] (> 336k samples have both height and BMI measured) as a replicate study. At *P* < 0.0008 (the Bonferroni threshold is calculated as 0.05/(32×2), since there are two traits), 27 out of the 32 SNPs were associated with both traits in the UK Biobank, including 17 of the 18 SNPs that were also associated with gene expression. Plots of −*log*_10_(*P*)-values for associations with height, BMI and expression in each tissue are presented in the Supplemental Figure 2 for the genomic regions containing the 27 replicated SNPs.

In *Supplemental Materials*, we also presented another set of analysis integrating GWAS summary statistics of Crohn’s disease and ulcerative colitis [36] with eQTL summary statistics from sigmoid colon (*n* = 318) and transverse colon (*n* = 368) tissues from GTEx. Both analyses showed that Primo can be used to detect SNPs with pleiotropic effects on (potentially more than two) complex traits while simultaneously providing mechanistic interpretations by examining their effects on cis-gene expression levels in trait-relevant tissue types. A majority of our detected and replicated pleiotropic SNPs do not have associations reaching genome-wide thresholds for both traits. Our analyses and results underscored the value of integrating GWAS summary statistics of multiple traits with eQTLs in relevant tissue types.

## Discussion

We proposed a general integrative genomics association approach – Primo – for assessing the joint associations across studies and data types, allowing for unknown study heterogeneity and sample correlation and taking only summary statistics as input. In elucidating the molecular mechanisms of trait-associated SNPs, we made a tailored development to conduct conditional association analysis in gene regions harboring known trait-associated SNPs and account for LD.

With the rapidly increasing availability of GWAS and omics QTL association summary statistics from different studies, populations, and cellular contexts, it is commonly observed that there could be multiple causal SNPs for different complex and omics traits in the same gene regions. Conducting integrative analysis of GWAS summary statistics and 1-2 sets of omics QTL statistics may provide only a partial view of the genomic activities in a region; meanwhile, if multiple omics QTL statistics are jointly analyzed, one also needs to consider the associations identified by chance and perform multiple testing adjustment. The advantage of Primo is that it can integrate a moderate to large number of sets of summary statistics from different data sources as input to provide a more comprehensive evaluation while also considering multiple testing adjustment. Additionally, Primo enjoys other unique advantages and shows great flexibility in integrative analysis. It allows the input summary statistics to be from independent, or partially overlapped studies with unknown study correlations. It detects SNPs with coordinated effects allowing different effect sizes (and different directions of effect sizes) on different types of traits. It can also integrate one-sided *P*-values if the same direction of effect sizes is expected and desired. Primo can identify SNPs in different combinations of association patterns to molecular omics and complex traits. Moreover, with the conditional association analysis of Primo, we can move one step beyond association towards causation by assessing whether a GWAS SNP is also an omics QTL while adjusting for the effects of multiple lead SNPs in a gene region. The conditional association analysis can reduce spurious omics-trait associations of GWAS SNPs due to LD with the lead omics SNPs.

We implemented two versions of Primo taking either *t*-statistics (or effect sizes and standard error estimates) or *P*-values as input. Primo is computationally very efficient and can analyze the joint associations of 30 million SNPs to five traits in dozens of minutes. We applied Primo to examine and interpret the associations to omics traits in tumor/normal tissues for known breast cancer susceptibility loci. We also applied Primo to integrate pairs of GWAS summary statistics of complex traits with eQTL summary statistics from trait-relevant tissue types from GTEx to detect pleiotropic effects and examine their mechanisms.

There are some caveats of the current work. First, our simulation results showed that when marginal study-specific sparsity parameters (*θ*’s) are over-specified, Primo may suffer from slightly inflated true FDR; whereas when those parameters are under-specified to an extent, there might not be much power loss. Therefore, we recommend a stringent specification of the marginal sparsity parameters, especially when there is limited *a priori* knowledge guiding the parameter specification. Second, there are many existing functional annotations for SNPs that are not incorporated in the current version of Primo but have also proved to be useful. We will explore this direction in future work. Last but not least, when jointly analyzing more than 15 sets of summary statistics, the computation time of Primo to assess all possible association patterns can increase substantially. The current work proposed a quick extension by applying Primo to groups of sets of summary statistics, while in a work-in-progress we will develop an integrative analysis method for jointly analyzing dozens of sets of summary statistics.

Primo is motivated by the analysis of trait-associated SNPs for their molecular trait-associations. It should be noted that Primo can also be broadly applied to many other settings when data integration is needed. Primo can be used to detect associations repeatedly observed in multiple correlated or independent conditions, and those repeatedly observed associations may enhance the confidence for new discoveries, or at least provide a more comprehensive examination of how those associations may occur in different conditions.

## Methods

### Estimating empirical null and alternative marginal density functions for each of the *J* studies using the limma method

For each of the *J* studies, we first adopt the limma method [49, 50] to calculate a set of *moderated t*-statistics by replacing the sample variance estimates in the classical *t*-statistic calculation with the posterior variances. Here we made a tailored development in the genetic association context by calculating the sample variance for each SNP based on the *t*-statistic and the minor allele frequency assuming that covariates are independent from genotypes. Alternatively, one may directly obtain the effect size estimate and its variance estimate as the summary statistics, if the information is available. The new variance shrinks the observed sample variance towards a prior that is estimated across all SNPs in the data, and stabilizes the variance estimation across the genome. It also penalizes the SNPs with large *t*-statistics but small variances.

Next, for each study *j*, we estimate the empirical null and alternative marginal density functions, 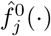 and 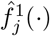, respectively, based on all the moderated *t*-statistics in the genome for the study. Here one needs to specify a key parameter for each study, the proportion of study-specific non-null statistics (i.e. with associations), 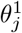. Note that we used 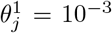 and 10^−5^ for omics QTL studies and GWAS, respectively, in the two applications. We then adopt the limma method to estimate 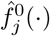 and 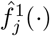 (illustrated in Figure 1D). Under the null hypothesis, the moderated *t*-statistic follows a *t*-distribution with a mean of zero and moderated degrees of freedom d_j_ in the *j*-th study, allowing for an empirical null distribution slightly deviating from the parametric *t*-distribution. Under the alternative, the moderated *t*-statistic follows a scaled *t*-distribution, still with degrees of freedom *d_j_* and a mean of zero allowing for different directions of effects in different studies, and a scaling factor *v_ij_* (*v_ij_* ≥ 1) estimated from the data. With the estimated marginal null and alternative density functions from each study, the joint density functions for all *K* association patterns can be calculated as described in the next subsection.

### Estimating pattern-specific multivariate density functions when input summary statistics are calculated from independent or overlapping samples

With *J* independent studies, the pattern-specific multivariate density function *D_k_* for the *k*-th association pattern is given by

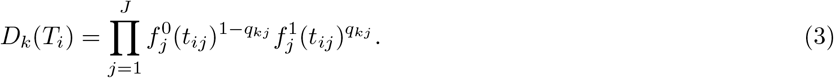

where *q_kj_* is the association status of the *k*-th pattern in study *j*. For example, given the association status being *q_k_* = (1, 1, 0, 0), the joint density *D_k_* is modeled as the product of the alternative marginal density functions from the first two studies and the null marginal density functions from the other two studies, 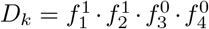.

In estimating a pattern-specific multivariate density function *D_k_* from *J* correlated studies, we obtain the empirical null and alternative marginal distributions as non-scaled and scaled *t*-distributions, respectively, in each of the *J* studies. Then we further approximate them with normal distributions with zero means and variances being 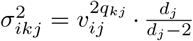, where *v_ij_* is the scaling factor under the alternative. Since *J* studies are correlated due to possible sample overlap with an unknown correlation matrix of **Γ**, similar to Urbut *et al.* (2019) [51] we pool all the statistics likely to be from the null pattern to estimate their correlation matrix as the estimate for **Γ**. Under certain assumptions, the correlation matrix of test statistics approximates the sample correlation matrix and the sample correlation under the null represents the correlation due to sample overlapping. Here we estimate the *J* × *J* correlation matrix using SNPs with absolute statistics less than 5 in all *J* studies. Then, we approximate the pattern-specific multivariate density function *D_k_* as 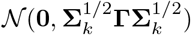, where **Σ**_*k*_ is a diagonal matrix with diagonal elements of 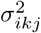’s. Primo separates sample correlations **Γ** from biological correlations/co-occurrences captured by *π_k_*’s in the subsequent estimation and inference.

### Conditional association analysis accounting for LD

To assess whether the trait-association of a SNP *i* reflects an independent causal variant or is simply due to being in LD with a nearby lead SNP *i′*, conditional association analysis is often conducted [52]. It tests the conditional association of SNP *i* with the trait of interest adjusting for the genotype of the lead SNP *i′* and other covariates. If SNP *i* is no longer statistically significant after adjusting for the lead SNP, it is unlikely that the trait-association of SNP *i* reflects an independent causal effect.

Following this idea, to assess whether a GWAS SNP is associated with omics traits due to it being in LD with lead omics QTLs, we propose to conduct conditional association analysis with summary statistics of the GWAS SNP and lead omics QTLs as input. Here we consider a GWAS SNP *i* of interest and a set of lead omics SNPs *I′* in the gene region, where *I′* = {1′,…, *L′*} is a set of indices. We can model the joint association statistics for SNPs *i* and *I′* in study *j*, i.e., (*t_ij_*, *t*_1′*j*_,…,*t_L′j_*), using a multivariate normal distribution, 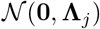, where **Λ**_*j*_ is the 1 + *L′* by 1 + *L′* variance-covariance matrix described as follows. The diagonal elements of **Λ**_*j*_ correspond to the study-specific variances of statistics of the SNPs. Specifically, the (1, 1) entry of **Λ**_*j*_ is given by 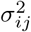, which is the marginal variance of the statistic *t_ij_* for SNP *i* in study *j* with 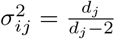 under the null and 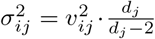 under the alternative. For each lead SNP *i′* ∈ *I′* with its most plausible association pattern *k_i′_*, the variance of the corresponding *t*-statistic *t_i′j_* is given by 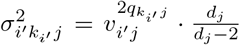. The off-diagonal elements of **Λ**_*j*_ are calculated based on the study-specific variances of the SNPs and the LD among the SNPs assuming additional covariates are independent of the SNP genotypes [53]. For instance, the covariance between *t_ij_* and *t_i′j_* is *σ_ij_* · *σ_i′k_i′_j_* · *ρ_ii′_* where 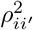 is the LD coefficient of the SNPs *i* and *i′*(∈ *I′*). Partitioning the variance-covariance matrix **Λ**_j_ as follows, 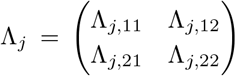 with sizes 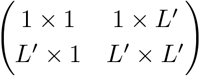, we can obtain the conditional null and alternative distributions for SNP *i* in study *j* as

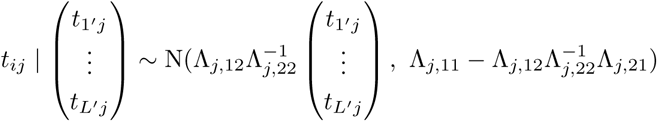

where 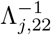 denotes the inverse of the matrix Λ_*j*,22_.

With the conditional null and alternative density functions for SNP *i* in study *j* adjusting for other lead omics SNPs in the region, we can proceed to obtain the pattern-specific *J*-variate density functions for all association patterns as outlined in the previous subsection and re-assess the probabilities of each association pattern in (1). We propose to conduct gene-level conditional association analysis accounting for LD structures only in selected gene regions, after the SNP-level association analysis.

### Primo for integrating P-values from multiple studies

In addition to integrating *t*-statistics or effect sizes and variance estimates, Primo can also jointly analyze *J* sets of *P*-values, chi-squared statistics, or other second-order association statistics. We model the pattern-specific multivariate density functions and still use equations (1) in obtaining the posterior probabilities for each SNP being in each pattern.

In estimating the marginal null and alternative density functions for each study *j*, 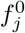 and 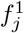 (as illustrated in Figure 1D), we make the following modification. We first take negative two times the log of *P*-values as our test statistics, **T**. Under the null hypothesis, *t_ij_* = −2log(*p_ij_*) follows a 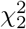 distribution. Under the alternative, *t_ij_*(*i* = 1,…, *m*) follows a mixture of non-central chi-squared distributions, which can be approximated by a scaled chi-squared distribution with certain degree of freedom, 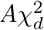 [54, 55]. To estimate a study-specific scaling factor *A_j_* > 0 and degree of freedom 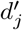 that best approximate the tail of the alternative distribution in study *j*, we use a numerical optimization algorithm to find values which minimize the differences between the *P*-values of *T_j_* under a mixture of 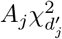 and 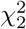 distributions given the mixing proportion 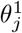 for the study, and their nominal *P*-values based on their ranks.

More specifically, let *t_ij_* = −2log(*p_ij_*) for SNP *i* in study *j*. Then the cumulative distribution function of *t_ij_* is given by

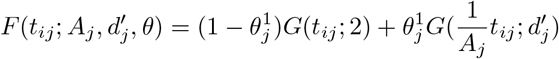

where *G*(·; *ν*) is the cumulative distribution function of a 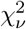 variable. Let *r_ij_* be the rank of SNP *i* in study *j* when the *t_ij_* are sorted in descending order. To estimate *A_j_* and 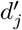, we use the optimization algorithms implemented in the R nloptr package: [56]

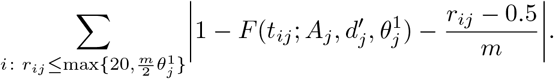

Since associations can be sparse (i.e., 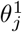 being close to zero) in the genome, it is more important to well approximate the tail of the alternative distribution than the first two moments (mean and variance). As such, we sum over the most extreme tail statistics or at least the 20 most extreme statistics. In *Supplemental Materials*, we have assessed the performance of the approximation via simulation studies, especially when associations are sparse. When the *J* studies are independent, the multivariate density function is modeled as the product of the individual density functions, as in Equation (3). When the *J* studies are correlated, we proceed in a similar manner as when *t*-statistics are used as input, except that the multivariate normal distribution is replaced by the multivariate gamma distribution.

### Extensions of Primo when *J* is large

When jointly analyzing a large number of sets of association summary statistics, the number of possible joint association patterns *K* = 2^J^ increases exponentially with the number of sets of statistics, *J*. When *J* = 15, there are 32,768 possible association patterns and the calculation for all *K* patterns can be computationally expensive. One may reduce the number of patterns under consideration to only the major and interpretable patterns [51]. However, the selection of major and interpretable patterns is still a challenge. Additional work is still needed in future research. When analyzing a large number of sets of association statistics of similar types (for example, integrating multiple sets of eQTLs from different GTEx tissue types for cross-tissue eQTLs), one possible strategy is to group sets of statistics into major and independent groups *g* = 1,…, *G*, each with *J_g_* < 10 sets of statistics. Then one can apply Primo to calculate the posterior probabilities within each group and take the products of the probabilities between groups to obtain the overall probabilities for all groups in the association patterns of interest. For example, the posterior probability of a SNP being associated with at least 1 (omics) trait in *G* groups of studies is given by

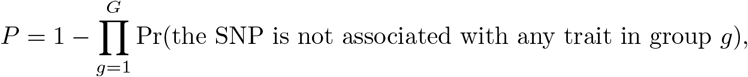

where the probability of the SNP being not associated with any trait in group *g* can be calculated by separately applying Primo to the low-dimensional *J_g_* set of statistics within the *g*-th group.

When jointly analyzing unbalanced numbers of summary statistics of different data types (e.g., 10 sets of eQTL and 1 set of pQTL statistics), caution should be taken as the joint association results can be dominated by one data type (here, eQTL), which is not ideal. One may first collapse those *J* sets of statistics by data types, and apply Primo in a hierarchical fashion to the (converted) summary statistics from multiple data types. This direction will be explored in future work.

### The connection of Primo to “colocalization” and meta-analysis methods

The Primo method shares some similarities with colocalization methods, as well as meta-analysis methods. Similar to colocalization methods [15, 16, 18, 17], Primo aims to integrate omics QTL with GWAS statistics to provide molecular mechanistic interpretations of known trait-associated SNPs. Additionally, the ‘coloc’[15] and ‘moloc’[16] methods assume that there is only up to one true causal variant in a region. However, the lead SNPs associated with expression levels in a gene can be different in different tissue types and cell types [6], and the SNPs for different omics QTLs may or may not share a same causal variant [21]. Motivated by those facts, Primo integrates GWAS statistics with omics and multi-omics QTL association statistics, and conducts conditional association analysis in gene regions harboring known trait-associated SNPs to assess their omics-trait associations accounting for LD with other lead SNPs for omics traits in the same gene regions.

Additionally, Primo enjoys a few advantages that are not shared with existing methods: Primo can integrate more than three sets of summary statistics; Primo requires only a total of *J* pre-specified parameters, 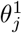’s (which often can be estimated from the data or based on a priori knowledge), and the results are not sensitive to under-specification of those parameters; Primo estimates the *π_k_*’s based on the data and separates the biological correlations/co-occurrences from sample correlations, i.e. allowing studies to be correlated. Additionally, Primo provides FDR estimates to guide the data-dependent choices of posterior probability cutoffs.

In comparison with meta-analysis, as a general association method Primo is more flexible in accounting for study heterogeneity, allowing different GWAS and omics QTL studies to have different effect sizes even in different directions. Note that if the same directions of effect sizes is expected for a biological reason, one can also use the one-side *P*-values as input in Primo. Primo does not require the samples to be independent among different studies, and can take summary statistics calculated from studies with independent, correlated, and/or overlapping samples. More importantly, in addition to the omibinus test in identifying associations in at least one study, Primo can identify SNPs in different combinations of association patterns, many of which may have biological interpretations.

Primo is a flexible integrative association method with only summary statistics as input. It makes minimal assumptions about the data structure underlying different sets of summary statistics, and assesses the joint associations across a moderate to large number of traits/data-types/conditions/studies.

## Abbreviations

GWAS: genome-wide association studies
SNP: single nucleotide polymorphism
QTL: quantitative trait locus
eQTL: expression quantitative trait locus
meQTL: methylation quantitative trait locus
pQTL: protein quantitative trait locus
Primo: Package in R for Integrative Multi-Omics association analysis
LD: linkage disequilibrium
GTEx: Genotype-Tissue Expression project
TCGA: The Cancer Genome Atlas
BCAC: Breast Canter Association Consortium
FDR: false discovery rate
BMI: body mass index

## Acknowledgements

We thank the GTEx Consortium. We thank Mr. Alvaro Barbeira, Mr. Rodrigo Bonazzola, Drs. Hae Kyung Im and Eric Gamazon for sharing GWAS summary statistics and providing valuable discussions, in particular for the analyses of pleiotropic effects.

## Availability of data and material

The R package Primo is available at https://github.com/kjgleason/primo.

## Author’s contributions

LCS and FY conceived the project. LSC, FY, and KJG developed the methods, performed the simulation studies, and wrote the manuscript. LSC and KJG analyzed the data. XH and BLP provided valuable suggestions to the development of the methods and analysis. KJG and FY developed the R software package.

## Ethics approval and consent to participate

Not applicable.

## Consent for publication

Not applicable.

## References

1. GWAS Catalog;. Available from: https://www.ebi.ac.uk/gwas/. Accessed: 31 Jan 2019.

2. MacArthur J, Bowler E, Cerezo M, Gil L, Hall P, Hastings E, et al. The new NHGRI-EBI Catalog of published genome-wide association studies (GWAS Catalog). Nucleic Acids Res. 2017 01;45(D1):D896–D901.

3. Edwards SL, Beesley J, French JD, Dunning AM. Beyond GWASs: illuminating the dark road from association to function. Am J Hum Genet. 2013 Nov;93(5):779–797.

4. HindorfF LA, Sethupathy P, Junkins HA, Ramos EM, Mehta JP, Collins FS, et al. Potential etiologic and functional implications of genome-wide association loci for human diseases and traits. Proc Natl Acad Sci USA. 2009 Jun;106(23):9362–9367.

5. Nicolae DL, Gamazon E, Zhang W, Duan S, Dolan ME, Cox NJ. Trait-associated SNPs are more likely to be eQTLs: annotation to enhance discovery from GWAS. PLoS Genet. 2010 Apr;6(4):e1000888.

6. The GTEx Consortium. Genetic effects on gene expression across human tissues. Nature. 2017 10;550(7675):204–213.

7. Johansson A, Enroth S, Palmblad M, Deelder AM, Bergquist J, Gyllensten U. Identification of genetic variants influencing the human plasma proteome. Proc Natl Acad Sci USA. 2013 Mar;110(12):4673–4678.

8. Smith AK, Kilaru V, Kocak M, Almli LM, Mercer KB, Ressler KJ, et al. Methylation quantitative trait loci (meQTLs) are consistently detected across ancestry, developmental stage, and tissue type. BMC Genomics. 2014 Feb;15:145.

9. McVicker G, van de Geijn B, Degner JF, Cain CE, Banovich NE, Raj A, et al. Identification of genetic variants that affect histone modifications in human cells. Science. 2013 Nov;342(6159):747–749.

10. Grubert F, Zaugg JB, Kasowski M, Ursu O, Spacek DV, Martin AR, et al. Genetic Control of Chromatin States in Humans Involves Local and Distal Chromosomal Interactions. Cell. 2015 Aug;162(5):1051–1065.

11. Li Y, van de Geijn B, Raj A, Knowles DA, Petti AA, Golan D, et al. RNA splicing is a primary link between genetic variation and diseas. Science. 2016;352(6285).

12. Pasaniuc B, Price AL. Dissecting the genetics of complex traits using summary association statistics. Nat Rev Genet. 2017 02;18(2):117–127.

13. Bonder MJ, Luijk R, Zhernakova DV, Moed M, Deelen P, Vermaat M, et al. Disease variants alter transcription factor levels and methylation of their binding sites. Nat Genet. 2017 01;49(1):131–138.

14. Suhre K, Arnold M, Bhagwat AM, Cotton RJ, Engelke R, Raffler J, et al. Connecting genetic risk to disease end points through the human blood plasma proteome. Nat Commun. 2017 02;8:14357.

15. Giambartolomei C, Vukcevic D, Schadt EE, Franke L, Hingorani AD, Wallace C, et al. Bayesian test for colocalisation between pairs of genetic association studies using summary statistics. PLoS Genet. 2014 May;10(5):e1004383.

16. Giambartolomei C, Zhenli Liu J, Zhang W, Hauberg M, Shi H, Boocock J, et al. A Bayesian Framework for Multiple Trait Colocalization from Summary Association Statistics. Bioinformatics. 2018 Mar;.

17. Wen X, Pique-Regi R, Luca F. Integrating molecular QTL data into genome-wide genetic association analysis: Probabilistic assessment of enrichment and colocalization [Journal Article]. PLoS Genet. 2017;13(3):e1006646. Available from: https://www.ncbi.nlm.nih.gov/pubmed/28278150.

18. Hormozdiari F, van de Bunt M, Segre AV, Li X, Joo JWJ, Bilow M, et al. Colocalization of GWAS and eQTL Signals Detects Target Genes. Am J Hum Genet. 2016 Dec;99(6):1245–1260.

19. Gusev A, Ko A, Shi H, Bhatia G, Chung W, Penninx BW, et al. Integrative approaches for large-scale transcriptome-wide association studies. Nat Genet. 2016 Mar;48(3):245–252.

20. Barbeira AN, Dickinson SP, Bonazzola R, Zheng J, Wheeler HE, Torres JM, et al. Exploring the phenotypic consequences of tissue specific gene expression variation inferred from GWAS summary statistics. Nat Commun. 2018 May;9(1):1825.

21. Pierce BL, Tong L, Argos M, Demanelis K, Jasmine F, Rakibuz-Zaman M, et al. Co-occurring expression and methylation QTLs allow detection of common causal variants and shared biological mechanisms. Nat Commun. 2018 02;9(1):804.

22. Battle A, Khan Z, Wang SH, Mitrano A, Ford MJ, Pritchard JK, et al. Genomic variation. Impact of regulatory variation from RNA to protein. Science. 2015 Feb;347(6222):664–667.

23. Chick JM, Munger SC, Simecek P, Huttlin EL, Choi K, Gatti DM, et al. Defining the consequences of genetic variation on a proteome-wide scale. Nature. 2016 06;534(7608):500–505.

24. Vandiedonck C. Genetic association of molecular traits: A help to identify causative variants in complex diseases. Clin Genet. 2018 Mar;93(3):520–532.

25. van der Wijst MGP, Brugge H, de Vries DH, Deelen P, Swertz MA, Franke L. Single-cell RNA sequencing identifies celltype-specific cis-eQTLs and co-expression QTLs. Nat Genet. 2018 Apr;50(4):493–497.

26. Chen L, Ge B, Casale FP, Vasquez L, Kwan T, Garrido-Martin D, et al. Genetic Drivers of Epigenetic and Transcriptional Variation in Human Immune Cells. Cell. 2016 11;167(5):1398–1414.

27. Yao C, Joehanes R, Johnson AD, Huan T, Esko T, Ying S, et al. Sex-and age-interacting eQTLs in human complex diseases. Hum Mol Genet. 2014 Apr;23(7):1947–1956.

28. Zhernakova DV, Deelen P, Vermaat M, van Iterson M, van Galen M, Arindrarto W, et al. Identification of context-dependent expression quantitative trait loci in whole blood. Nat Genet. 2017 01;49(1):139–145.

29. Sivakumaran S, Agakov F, Theodoratou E, Prendergast JG, Zgaga L, Manolio T, et al. Abundant pleiotropy in human complex diseases and traits. Am J Hum Genet. 2011 Nov;89(5):607–618.

30. Pickrell JK, Berisa T, Liu JZ, Segurel L, Tung JY, Hinds DA. Detection and interpretation of shared genetic influences on 42 human traits. Nat Genet. 2016 07;48(7):709–717.

31. Parkes M, Cortes A, van Heel DA, Brown MA. Genetic insights into common pathways and complex relationships among immune-mediated diseases. Nat Rev Genet. 2013 Sep;14(9):661–673.

32. Cross-Disorder Group of the Psychiatric Genomics Consortium. Identification of risk loci with shared effects on five major psychiatric disorders: a genome-wide analysis. Lancet. 2013 Apr;381(9875):1371–1379.

33. Wu YH, Graff RE, Passarelli MN, Hoffman JD, Ziv E, Hoffmann TJ, et al. Identification of Pleiotropic Cancer Susceptibility Variants from Genome-Wide Association Studies Reveals Functional Characteristics. Cancer Epidemiol Biomarkers Prev. 2018 Jan;27(1):75–85.

34. The Cancer Genome Atlas Network. Comprehensive molecular portraits of human breast tumours. Nature. 2012;490:61–70.

35. Michailidou K, Lindstrom S, Dennis J, Beesley J, Hui S, Kar S, et al. Association analysis identifies 65 new breast cancer risk loci. Nature. 2017 Nov;551(7678):92–94.

36. Liu JZ, van Sommeren S, Huang H, Ng SC, Alberts R, Takahashi A, et al. Association analyses identify 38 susceptibility loci for inflammatory bowel disease and highlight shared genetic risk across populations. Nat Genet. 2015 Sep;47(9):979–986.

37. Wood AR, Esko T, Yang J, Vedantam S, Pers TH, Gustafsson S, et al. Defining the role of common variation in the genomic and biological architecture of adult human height. Nat Genet. 2014 Nov;46(11):1173–1186.

38. Locke AE, Kahali B, Berndt SI, Justice AE, Pers TH, Day FR, et al. Genetic studies of body mass index yield new insights for obesity biology. Nature. 2015 Feb;518(7538):197–206.

39. Churchhouse C, Neale B. Rapid GWAS of thousands of phenotypes for 337,000 samples in the UK Biobank; 2017. Available from: http://www.nealelab.is/blog/2017/7719/rapid-gwas-of-thousands-of-phenotypes-for-337000-samples-in-the-uk-biobank.

40. Wei Y, Tenzen Y, Ji H. Joint analysis of differential gene expression in multiple studies using correlation motifs. Biostatistics. 2015;16(1):31–46.

41. Dempster AP, Laird NM, Rubin DB. Maximum Likelihood from Incomplete Data via the EM Algorithm. JRSS, Series B. 1977;39(1):1–38.

42. Storey JD, Tibshirani R. Statistical significance for genomewide studies. Proc Natl Acad Sci USA. 2003 Aug;100(16):9440–9445.

43. Fisher RA. The Correlation between Relatives on the Supposition of Mendelian Inheritance. Philosophical Transactions of the Royal Society of Edinburgh. 1918 Apr;52:399–433.

44. Storey JD, Bass AJ, Dabney A, Robinson D. qvalue: Q-value estimation for false discovery rate control; 2015. Available from: http://github.com/jdstorey/qvalue.

45. Guo X, Lin W, Bao J, Cai Q, Pan X, Bai M, et al. A Comprehensive cis-eQTL Analysis Revealed Target Genes in Breast Cancer Susceptibility Loci Identified in Genome-wide Association Studies. Am J Hum Genet. 2018 05;102(5):890–903.

46. Pai AA, Pritchard JK, Gilad Y. The genetic and mechanistic basis for variation in gene regulation. PLoS Genet. 2015 Jan;11(1):e1004857.

47. Mertins P, Mani DR, Ruggles KV, Gillette MA, Clauser KR, Wang P, et al. Proteogenomics connects somatic mutations to signalling in breast cancer. Nature. 2016 06;534(7605):55–62.

48. Solovieff N, Cotsapas C, Lee PH, Purcell SM, Smoller JW. Pleiotropy in complex traits: challenges and strategies. Nat Rev Genet. 2013 Jul;14(7):483–495.

49. Smyth GK. Linear models and empirical bayes methods for assessing differential expression in microarray experiments. Stat Appl Genet Mol Biol. 2004;3:Article3.

50. Ritchie ME, Phipson B, Wu D, Hu Y, Law CW, Shi W, et al. limma powers differential expression analyses for RNA-sequencing and microarray studies. Nucleic Acids Research. 2015;43(7):e47.

51. Urbut SM, Wang G, Carbonetto P, Stephens M. Flexible statistical methods for estimating and testing effects in genomic studies with multiple conditions. Nature Genetics. 2019;51:187–195.

52. Schaid DJ, Chen W, Larson NB. From genome-wide associations to candidate causal variants by statistical fine-mapping. Nature Review Genetics. 2-18;p. 491–504.

53. Yang J, Ferreira T, Morris AP, Medland SE, Madden PA, Heath AC, et al. Conditional and joint multiple-SNP analysis of GWAS summary statistics identifies additional variants influencing complex traits. Nat Genet. 2012 Mar;44(4):369–375.

54. Satterthwaite FE. An Approximate Distribution of Estimates of Variance Components. Biometrics. 1946;2:110–114.

55. Solomon H, Stephens MA. Distribution of a Sum of Weighted Chi-Square Variables. J Amer Statist Assoc. 1977;72:881–885.

56. Johnson SG. The NLopt nonlinear-optimization package; 2018. Available from: http://ab-initio.mit.edu/nlopt.

