## Supplemental Materials for "Primo: integration of multiple GWAS and omics QTL summary statistics for elucidation of molecular mechanisms of trait-associated SNPs and detection of pleiotropy in complex traits"

METHOD

### Supplemental Methods

#### Genotyping and imputation

GTEx samples underwent Whole Genome Sequencing at an average coverage of 30X on Illumina HiSeq 2000 or Illumina HiSeq X. Additional details about the genotyping pipeline and sample and variant quality control have been reported elsewhere [1].

Genotype data and clinical covariates for TCGA breast cancer subjects were downloaded through the Genomic Data Commons (GDC) Data Portal [2]. TCGA subjects were genotyped on the Affymetrix Genome-wide Human SNP Array 6.0. Germline genotypes were measured in blood-derived DNA samples primarily. For subjects missing genotyping from blood samples, genotype measured in solid normal tissue was used as a surrogate. Restricting to bi-allelic variants on autosomes yielded 859,193 SNPs. After removing subjects with duplicate blood genotypes that did not match (i.e. labeling problems), there were 1094 subjects remaining. IMPUTE2 [3] was used to conduct genotype imputation using 1000 Genomes as the reference panel (phase3 v5) [4]. We performed 30 MCMC iterations, discarding the first 10 as burn-in, using 1 MB intervals for inference. SNPs with an imputation info score $<$ 0.3 or with a minor allele frequency (MAF) $<$ 0.01 were removed post-imputation.

#### Gene expression

GTEx RNA sequencing was performed using the Illumina TrueSeq RNA Sequencing platform. Data was aligned using STAR (v2.5.3a) [5]. Picard [6] was used to mark and remove duplicate reads. Transcripts were quantified using RSEM [7]. RNA-SeQC [8] was used for quality control and gene-level expression quantification, and TMM [9] was used to normalize read counts. Additional details on the RNA-Sequencing pipeline and processing are reported elsewhere [1].

Gene expression, protein abundance, and DNA methylation data for TCGA subjects were downloaded using TCGA-Assembler 2 [10]. RNA sequencing was performed in tumor tissues using the Illumina HiSeq 2000 RNA Sequencing platform. RNA-Seq expression levels were estimated using Fragments per Kilobase of transcript per Million mapped reads (FPKM) and mapped to each gene using HTSeq-count [11]. Expression data were quantile normalized prior to analysis.

#### DNA methylation

DNA methylation was measured in tumor tissue samples using the Infinium HumanMethylation450 BeadChip. The level of methylation at each CpG site was measured as a $\beta$ value ranging from 0 (completely unmethylated) to 1 (completely methylated). Data were quantile normalized prior to analysis.

#### Protein abundance

Protein abundance was measured in tumor tissue samples using iTRAQ (isobaric Tag for relative and Absolute Quantitation) mass-spectometry (MS) in experiments conducted by the Clinical Proteomic Tumor Analysis Consortium (CPTAC) [12]. The protein abundance was measured using the Log Ratio (i.e. the log of the ratio between the spectral count of a protein in a sample versus the spectral count of the protein in the reference sample). We restricted our analysis of protein abundance to the 77 high-quality samples as identified by Mertins, *et al.* [13]. We further restricted the analysis to the 74 of these 77 samples from female subjects with measured tumor purity scores [14].

#### QTL analyses

GTEx *cis*-eQTL summary statistics were generated by linear regression as implemented in FastQTL [15], adjusting for sex, genotyping platform, five genotype principal components (PCs), and up to 60 PEER [16] variables.

Prior to QTL analyses, expression, methylation and protein measurements for TCGA Samples were transformed to the quantiles of the standard normal distribution (separately for each gene, CpG site or protein). Imputed genotypes were converted to dosage format (i.e. expected counts of the reference allele). We generated *cis*-QTL summary statistics using linear regression as implemented in Matrix eQTL [17], adjusting for subject covariates. QTL analyses were restricted to female subjects. Covariates included tumor purity scores [14], cancer stage, histological subtype (infiltrating ductal, infiltrating lobular, mucinous, metaplastic, mixed histology or other), estrogen receptor (ER) and progesterone receptor (PR) status, and genotype principal components (15 PCs for expression and methylation; 3 PCs for protein). Genotype PCs were generated in PLINK 1.90 [18, 19] using measured bi-allelic variants on autosomes with the following filters: minor allele frequency $\geq$ 0.05; Hardy-Weinberg Equilibrium p-value $\geq 0.0001$; pairwise linkage disequilibrium $R^2 \leq 0.2$.

*Cis* associations were defined as: $<$ 250 kb apart from transcription start site for expression and protein; $<$ 50 kb apart from a CpG site for methylation. For GWAS-reported SNPs that were missing based on these criteria, we selected the nearest gene (based on distance between the SNP and transcription start site) for which both gene expression and protein abundance levels were available (Application I in the main text) or for which gene expression levels were measured in both tissues (Application II in the main text). For these regions, we included associations $<$ 1 Mb apart from the transcription start site.

### Supplemental Results

#### Simulation of $\pi_k$ estimation for mis-specification of alternative proportions

We evaluated the performance of estimating the proportion of SNPs belonging to each association pattern ($\pi_k$) when the marginal alternative proportions $\theta_j^1$'s are

| Scenario | Specific. | Method | $\pi_k(\%)$ | | | | | | | |
|---|---|---|---|---|---|---|---|---|---|---|
| | | | $q_k$=(0 0 0) | (1 0 0) | (0 1 0) | (0 0 1) | (1 1 0) | (1 0 1) | (0 1 1) | (1 1 1) |
| S1a | | | Independent | | | | | | | |
| | | True | 99.720 | 0.070 | 0.070 | 0.070 | 0.020 | 0.020 | 0.020 | 0.010 |
| | Under | Primo ($t$) | 99.767 | 0.061 | 0.061 | 0.061 | 0.015 | 0.015 | 0.015 | 0.006 |
| | | Primo ($P$) | 99.822 | 0.049 | 0.049 | 0.049 | 0.009 | 0.009 | 0.009 | 0.002 |
| | Over | Primo ($t$) | 99.648 | 0.091 | 0.091 | 0.091 | 0.022 | 0.022 | 0.022 | 0.014 |
| | | Primo ($P$) | 99.424 | 0.158 | 0.158 | 0.157 | 0.028 | 0.028 | 0.028 | 0.018 |
| | | | Correlated | | | | | | | |
| | | True | 99.720 | 0.070 | 0.070 | 0.070 | 0.020 | 0.020 | 0.020 | 0.010 |
| | Under | Primo ($t$) | 99.766 | 0.061 | 0.061 | 0.061 | 0.015 | 0.015 | 0.015 | 0.007 |
| | | Primo ($P$) | 99.823 | 0.048 | 0.048 | 0.048 | 0.010 | 0.010 | 0.010 | 0.003 |
| | Over | Primo ($t$) | 99.648 | 0.091 | 0.091 | 0.091 | 0.022 | 0.022 | 0.022 | 0.014 |
| | | Primo ($P$) | 99.436 | 0.147 | 0.147 | 0.146 | 0.035 | 0.035 | 0.035 | 0.021 |
| S1b | | | Independent | | | | | | | |
| | | True | 99.840 | 0.070 | 0.070 | 0.0007 | 0.020 | 0.0002 | 0.0002 | 0.0001 |
| | Under | Primo ($t$) | 99.868 | 0.058 | 0.058 | 0.0024 | 0.014 | 0.0001 | 0.0002 | 0.0001 |
| | | Primo ($P$) | 99.903 | 0.044 | 0.044 | 0.0004 | 0.008 | 0.0001 | 0.0001 | 0.0001 |
| | Over | Primo ($t$) | 99.785 | 0.091 | 0.091 | 0.0091 | 0.023 | 0.0004 | 0.0004 | 0.0002 |
| | | Primo ($P$) | 99.634 | 0.160 | 0.160 | 0.0147 | 0.030 | 0.0005 | 0.0005 | 0.0002 |
| | | | Correlated | | | | | | | |
| | | True | 99.840 | 0.070 | 0.070 | 0.0007 | 0.020 | 0.0002 | 0.0002 | 0.0001 |
| | Under | Primo ($t$) | 99.867 | 0.058 | 0.058 | 0.0021 | 0.014 | 0.0002 | 0.0002 | 0.0001 |
| | | Primo ($P$) | 99.903 | 0.044 | 0.044 | 0.0004 | 0.008 | 0.0001 | 0.0001 | 0.0001 |
| | Over | Primo ($t$) | 99.779 | 0.093 | 0.093 | 0.0114 | 0.022 | 0.0005 | 0.0005 | 0.0002 |
| | | Primo ($P$) | 99.640 | 0.153 | 0.153 | 0.0133 | 0.034 | 0.0019 | 0.0019 | 0.0020 |

**Table S1 Average estimates of $\hat{\pi}$ under mis-specification of the alternative proportions, $\theta_j^1$'s. In the simulations, $\hat{\theta}_j^1 = \theta_j^1/10$ when underspecified and $\hat{\theta}_j^1 = \theta_j^1 \times 2$ when overspecified. Scenario S1a simulates sparse associations for $J = 3$ traits. Scenario S1b simulates even sparser associations for the third trait.**

mis-specified. Results are shown in Table S1. Scenario S1a simulates sparse associations, with true ($\pi_k = 7 \times 10^{-4}$, $2 \times 10^{-4}$, $1 \times 10^{-4}$) for SNPs being associated with only one, exactly two, and all three traits, respectively. Scenario S1b simulated even sparser associations for the third trait, with $\pi_k = (7 \times 10^{-6}, 2 \times 10^{-6}, 1 \times 10^{-6})$ for SNPs being associated with only the third, the third and first or second, and all three traits, respectively. The $\theta_j^1$'s are under-specified as $\theta_j^1/10$ and over-specified as $\theta_j^1 \times 2$. Primo estimates the $\pi_k$'s with reasonable accuracy even when the associations are very sparse and when the marginal alternative proportions $\theta_j^1$'s are mis-specified.

### Numerical optimization simulation for p-values

We evaluated the performance of Primo in estimating the scaling factor ($A_j$) and degrees of freedom ($d_j'$) parameters of the alternative distribution for $t_{ij} = -2\log(p_{ij})$ $(1,\ldots,m)$. Under different specifications of the proportion of statistics coming from the alternative distribution ($\theta_j^1$), we simulated 10 million test statistics. Test statistics under the null hypothesis of no association were simulated from a $\chi_2^2$ distribution; test statistics under the alternative were simulated from a $A_j\chi^2_{d_j'}$ distribution. Figure S1 compares the density curves of the true alternative density to the alternative densities estimated by Primo over 1000 simulations for $A_j = 4.5$ and $d_j' = 7$. As shown, the density curves estimated by Primo reasonably approximate the true density curve even when the proportion of statistics coming from the alternative distribution are sparse ($\theta_j^1 = 1 \times 10^{-4}$ ; panel S1A) or very sparse ($\theta_j^1 = 1 \times 10^{-5}$ ; panel S1B).

### Detecting SNPs with pleiotropic effects on Crohn's disease and ulcerative colitis and elucidating their mechanisms

Here we applied Primo to integrate GWAS summary statistics of Crohn's disease and ulcerative colitis from a study of over 20,000 samples of European Ancestry conducted by the International Inflammatory Bowel Disease (IBD) Genetics Consortium [20] with eQTL summary statistics from sigmoid colon ($n = 318$) and transverse colon ($n = 368$) tissues from GTEx. Of the 232 SNPs reported in the initial meta-analysis, 68 have reached genome-wide significance for at least one of Crohn's disease or ulcerative colitis in the genome-wide analysis, and among those, 67 SNPs could be mapped to GTEx SNPs in cis with at least one gene measured in each tissue. At the 80% probability cutoff (estimated FDR of 0.8%, 5.6% and 6.4%) and after conditional association analysis accounting for LD, 37, 15 and 11 of the 67 SNPs were associated with both complex traits, both complex traits plus gene expression in at least 1 tissue, and both complex traits plus gene expression in both tissues, respectively. We used GWAS summary statistics of self-reported Crohn's disease and self-reported ulcerative colitis from the UK Biobank [21] to replicate our findings. At $P < 0.0007$ (0.05/37/2), 4 of the 37 SNPs were associated with both traits in the UK Biobank. The relatively low replication rate might be due to the fact that in the UK Biobank data, self-reported disease status was used for both traits and the numbers of cases of Crohn's diseases and ulcerative colitis are low (1,045 and 1,787, respectively).

### Reference

**Author details**

[1]Department of Public Health Sciences, University of Chicago, 5841 South Maryland Ave MC2000, Chicago, IL 60637. [2]Department of Biostatistics and Informatics, Colorado School of Public Health, University of Colorado Denver, 13001 E. 17th Place, Aurora, Colorado 80045. [3]Department of Human Genetics, University of Chicago, 920 E 58th St, Chicago, IL 60637.

**References**

1. The GTEx Consortium. Genetic effects on gene expression across human tissues. Nature. 2017 10;550(7675):204–213.
2. Grossman RL, Heath AP, Ferretti V, Varmus HE, Lowy DR, Kibbe WA, et al. Toward a Shared Vision for Cancer Genomic Data. N Engl J Med. 2016 Sep;375(12):1109–1112.
3. Howie BN, Donnelly P, Marchini J. A flexible and accurate genotype imputation method for the next generation of genome-wide association studies. PLoS Genet. 2009 Jun;5(6):e1000529.
4. Auton A, Brooks LD, Durbin RM, Garrison EP, Kang HM, Korbel JO, et al. A global reference for human genetic variation. Nature. 2015 Oct;526(7571):68–74.
5. Dobin A, Davis CA, Schlesinger F, Drenkow J, Zaleski C, Jha S, et al. STAR: ultrafast universal RNA-seq aligner. Bioinformatics. 2013 Jan;29(1):15–21.
6. Broad Institute. Picard Tools; 2019. Available from: http://broadinstitute.github.io/picard/.
7. Li B, Dewey CN. RSEM: accurate transcript quantification from RNA-Seq data with or without a reference genome. BMC Bioinformatics. 2011 Aug;12:323.
8. DeLuca DS, Levin JZ, Sivachenko A, Fennell T, Nazaire MD, Williams C, et al. RNA-SeQC: RNA-seq metrics for quality control and process optimization. Bioinformatics. 2012 Jun;28(11):1530–1532.
9. Robinson MD, Oshlack A. A scaling normalization method for differential expression analysis of RNA-seq data. Genome Biol. 2010;11(3):R25.
10. Wei L, Jin Z, Yang S, Xu Y, Zhu Y, Ji Y. TCGA-assembler 2: software pipeline for retrieval and processing of TCGA/CPTAC data. Bioinformatics. 2018 May;34(9):1615–1617.
11. Anders S, Pyl PT, Huber W. HTSeq–a Python framework to work with high-throughput sequencing data. Bioinformatics. 2015 Jan;31(2):166–169.
12. Ellis MJ, Gillette M, Carr SA, Paulovich AG, Smith RD, Rodland KK, et al. Connecting genomic alterations to cancer biology with proteomics: the NCI Clinical Proteomic Tumor Analysis Consortium. Cancer Discov. 2013 Oct;3(10):1108–1112.
13. Mertins P, Mani DR, Ruggles KV, Gillette MA, Clauser KR, Wang P, et al. Proteogenomics connects somatic mutations to signalling in breast cancer. Nature. 2016 06;534(7605):55–62.
14. Aran D, Sirota M, Butte AJ. Systematic pan-cancer analysis of tumour purity. Nat Commun. 2015 Dec;6:8971.
15. Ongen H, Buil A, Brown AA, Dermitzakis ET, Delaneau O. Fast and efficient QTL mapper for thousands of molecular phenotypes. Bioinformatics. 2016 05;32(10):1479–1485.
16. Stegle O, Parts L, Piipari M, Winn J, Durbin R. Using probabilistic estimation of expression residuals (PEER) to obtain increased power and interpretability of gene expression analyses. Nat Protoc. 2012 Feb;7(3):500–507.
17. Shabalin AA. Matrix eQTL: ultra fast eQTL analysis via large matrix operations. Bioinformatics. 2012 May;28(10):1353–1358.
18. Chang CC, Chow CC, Tellier LC, Vattikuti S, Purcell SM, Lee JJ. Second-generation PLINK: rising to the challenge of larger and richer datasets. Gigascience. 2015;4:7.
19. Purcell S, Chang C. PLINK 1.90; 2017. Available from: http://www.cog-genomics.org/plink/1.9/.

20. Liu JZ, van Sommeren S, Huang H, Ng SC, Alberts R, Takahashi A, et al. Association analyses identify 38 susceptibility loci for inflammatory bowel disease and highlight shared genetic risk across populations. Nat Genet. 2015 Sep;47(9):979–986.
21. Churchhouse C, Neale B. Rapid GWAS of thousands of phenotypes for 337,000 samples in the UK Biobank; 2017. Available from: http://www.nealelab.is/blog/2017/7/19/rapid-gwas-of-thousands-of-phenotypes-for-337000-samples-in-the-uk-biobank.

### Supplementary figures

**Figure S1 Primo reasonably estimates the scaling factor ($A_j$) and degrees of freedom ($d'_j$) for the alternative distribution, even when associations are sparse.** For $A_j = 4.5$ and $d'_j = 7$, the true density curve is shown by the dotted red curve in A and B. For $\theta_j^1 = 1 \times 10^{-4}$ (A) and $\theta_j^1 = 1 \times 10^{-5}$ (B), the density curves estimated by the median estimates of the parameters over 1000 simulations are given by the black curve. The shaded gray area shows the curves of the parameters between the $5th$ and $95th$ percentiles.

**Figure S2 Locus-zoom plots of $-\log_{10}(P)$-values for associations with Height, BMI (from GWAS) and gene expression levels (from GTEx eQTL analysis) for gene regions with pleiotropic SNPs being replicated.** Each page shows a set of four association plots for a locus: one for Height, one for BMI, and one for gene expression in each of the two tissue types – subcutaneous adipose and skeletal muscle. In each plot, the GWAS-reported SNP is colored red if associated with the given trait and colored blue if not associated with the given trait (at an 80% posterior probability threshold and after conditional association analysis accounting for LD). Lead omics SNPs which were adjusted for in conditional association analysis are shown by a green "X". For any SNP which was associated with expression of multiple genes, the gene with which it has the strongest association is presented in each tissue.

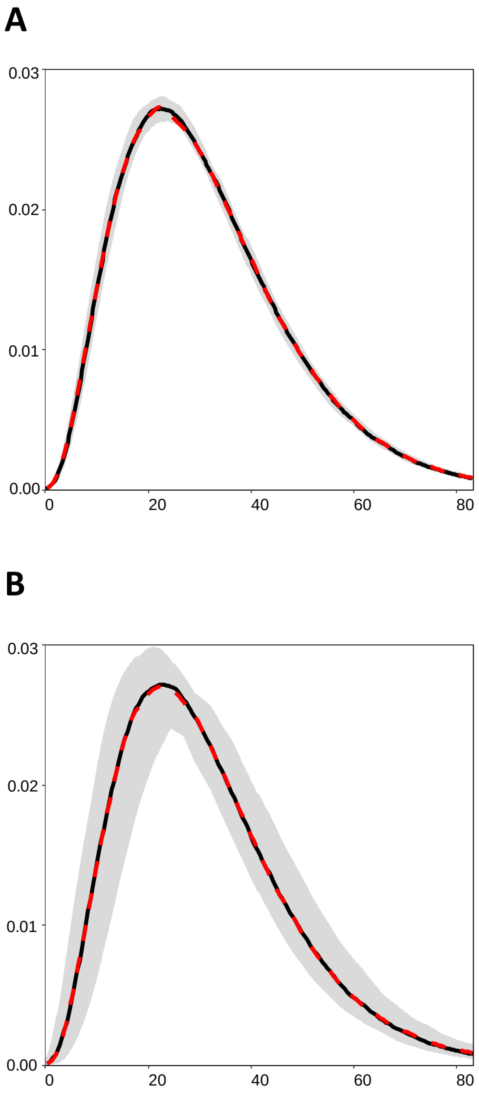
A
0.03
0.02
0.01
0.00
0
20
40
60
80
B
0.03
0.02
0.01
0.00
0
20
40
60
80

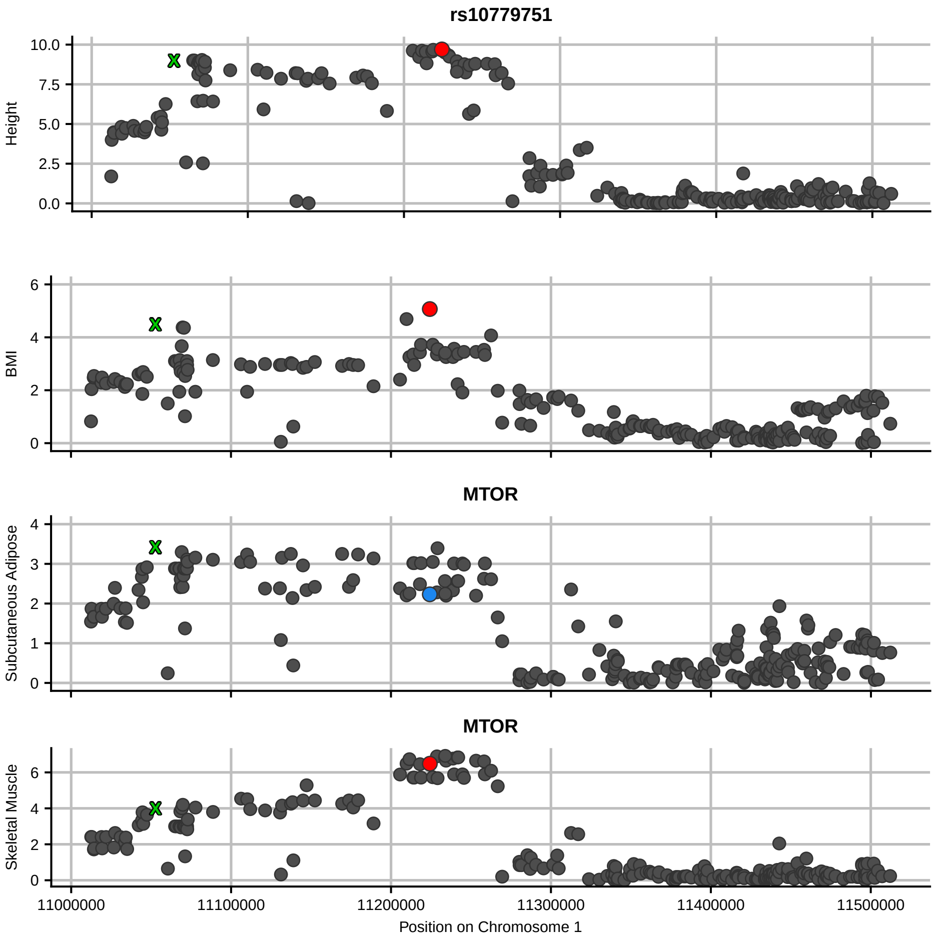

rs10779751
Height
10.0
7.5
5.0
2.5
0.0
BMI
6
4
2
0
MTOR
Subcutaneous Adipose
4
3
2
1
0
MTOR
Skeletal Muscle
6
4
2
0
11000000
11100000
11200000
11300000
11400000
11500000
Position on Chromosome 1

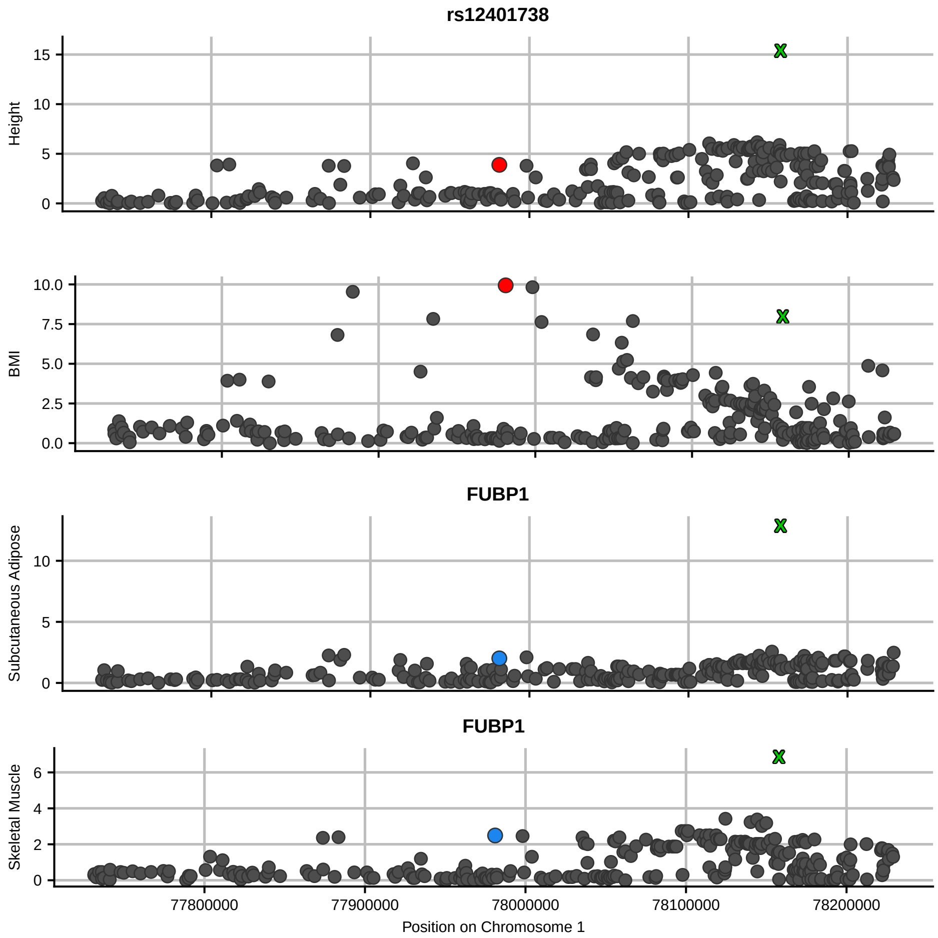

rs12401738
Height
BMI
FUBP1
Subcutaneous Adipose
FUBP1
Skeletal Muscle
77800000
77900000
78000000
78100000
78200000
Position on Chromosome 1

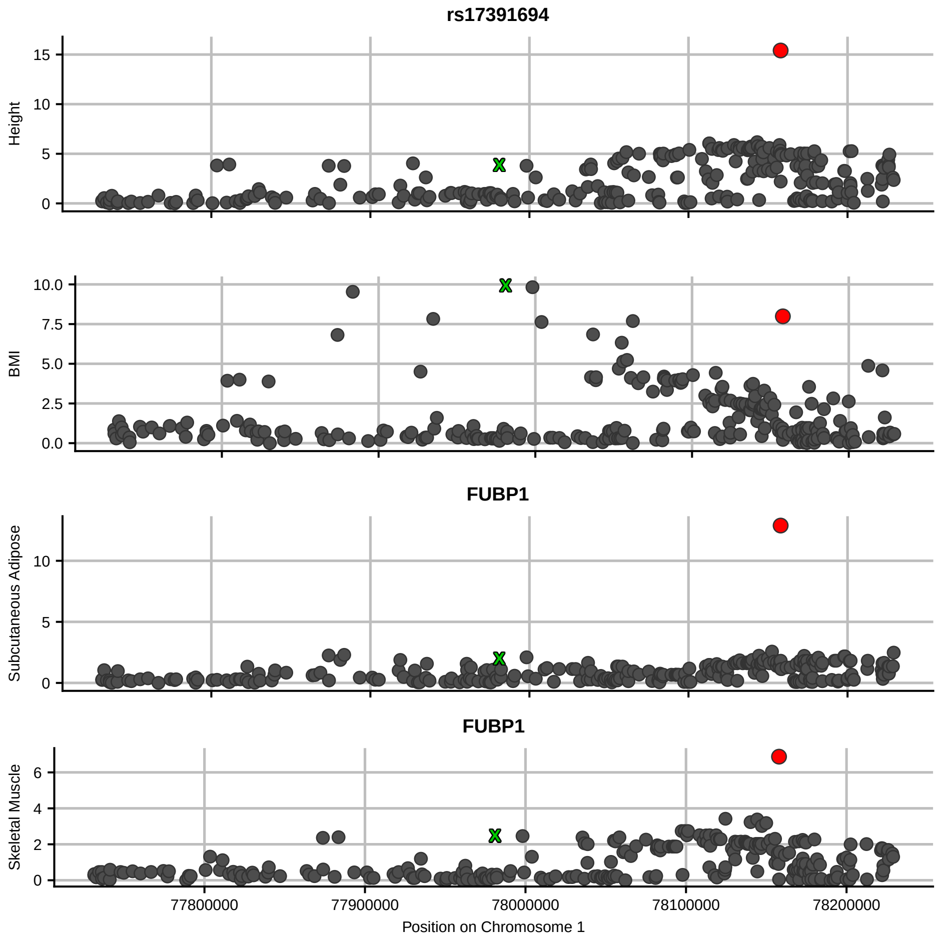

rs17391694
Height
BMI
FUBP1
Subcutaneous Adipose
FUBP1
Skeletal Muscle
77800000
77900000
78000000
78100000
78200000
Position on Chromosome 1

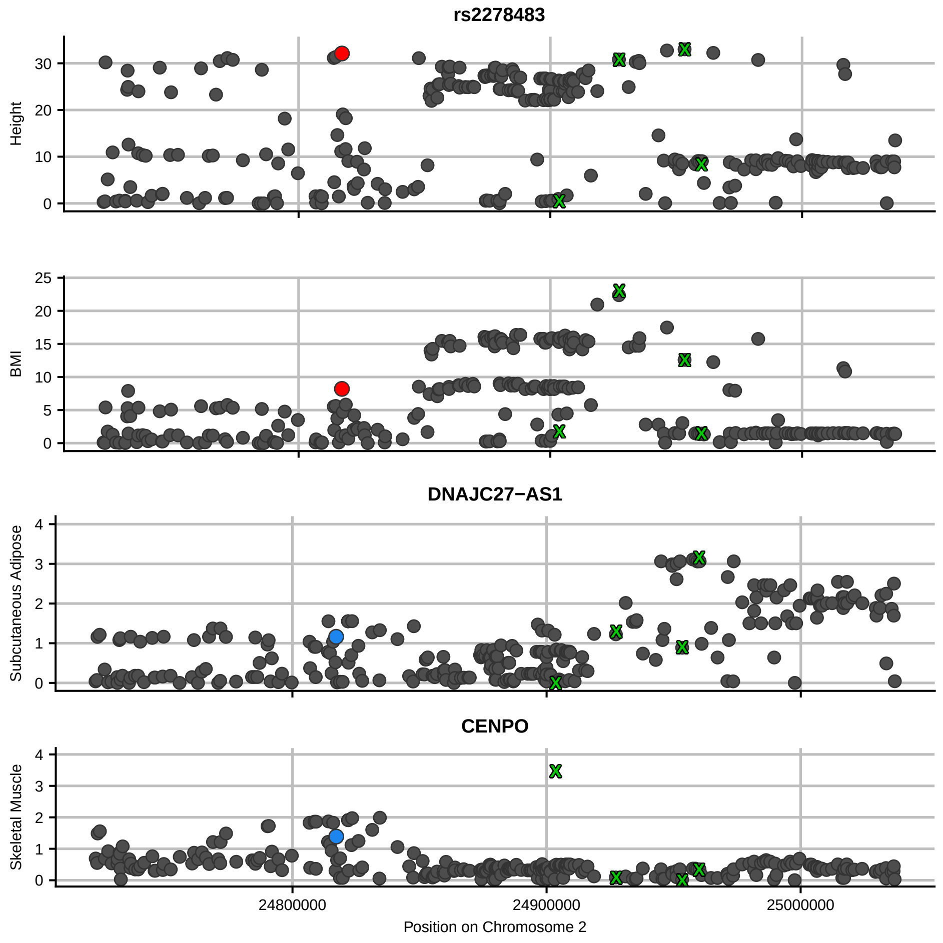

rs2278483
Height
BMI
DNAJC27−AS1
Subcutaneous Adipose
CENPO
Skeletal Muscle
24800000
24900000
25000000
Position on Chromosome 2

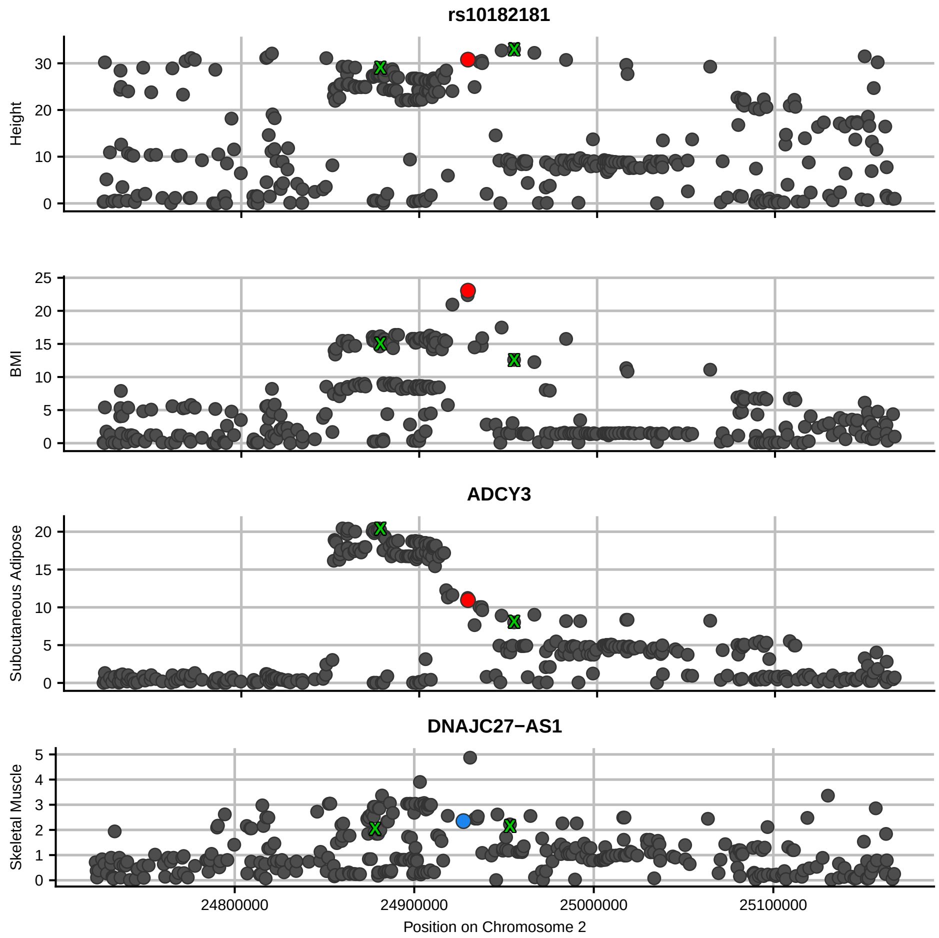

rs10182181
Height
BMI
ADCY3
Subcutaneous Adipose
DNAJC27−AS1
Skeletal Muscle
24800000
24900000
25000000
25100000
Position on Chromosome 2

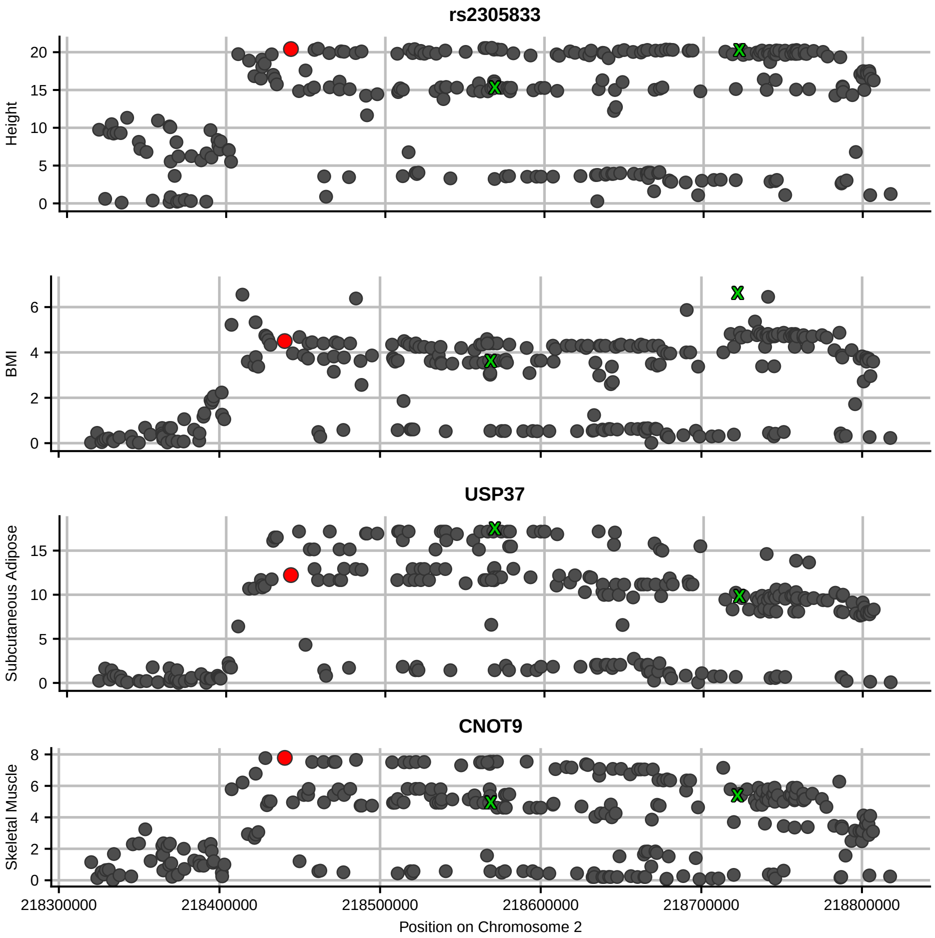

rs2305833
Height
BMI
USP37
Subcutaneous Adipose
CNOT9
Skeletal Muscle
218300000
218400000
218500000
218600000
218700000
218800000
Position on Chromosome 2

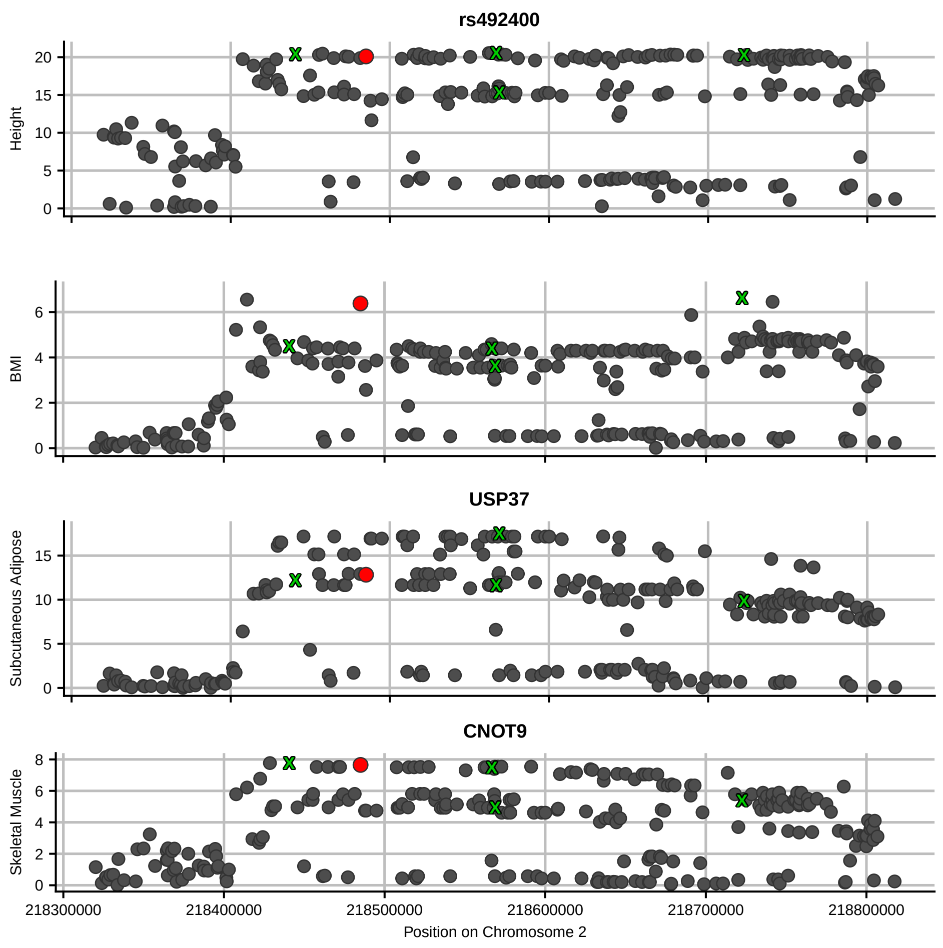
rs492400
Height
BMI
USP37
Subcutaneous Adipose
CNOT9
Skeletal Muscle
218300000
218400000
218500000
218600000
218700000
218800000
Position on Chromosome 2

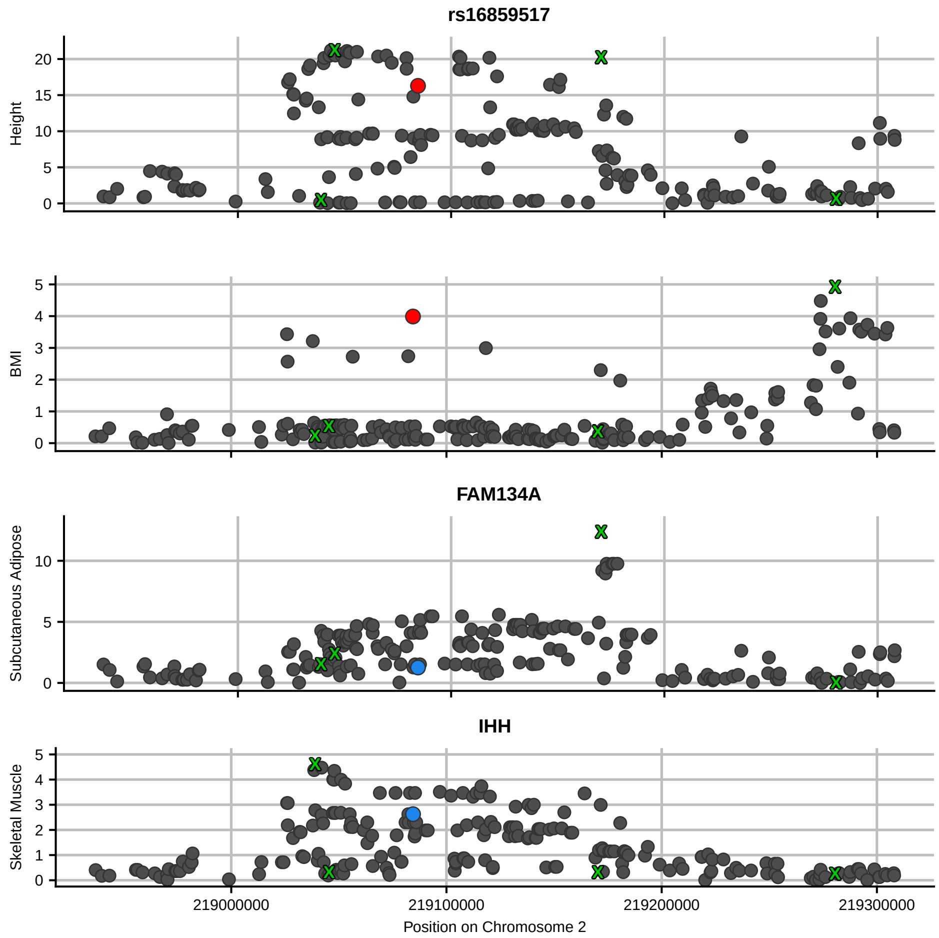

rs16859517
Height
20
15
10
5
0
BMI
5
4
3
2
1
0
FAM134A
Subcutaneous Adipose
10
5
0
IHH
Skeletal Muscle
5
4
3
2
1
0
219000000
219100000
219200000
219300000
Position on Chromosome 2

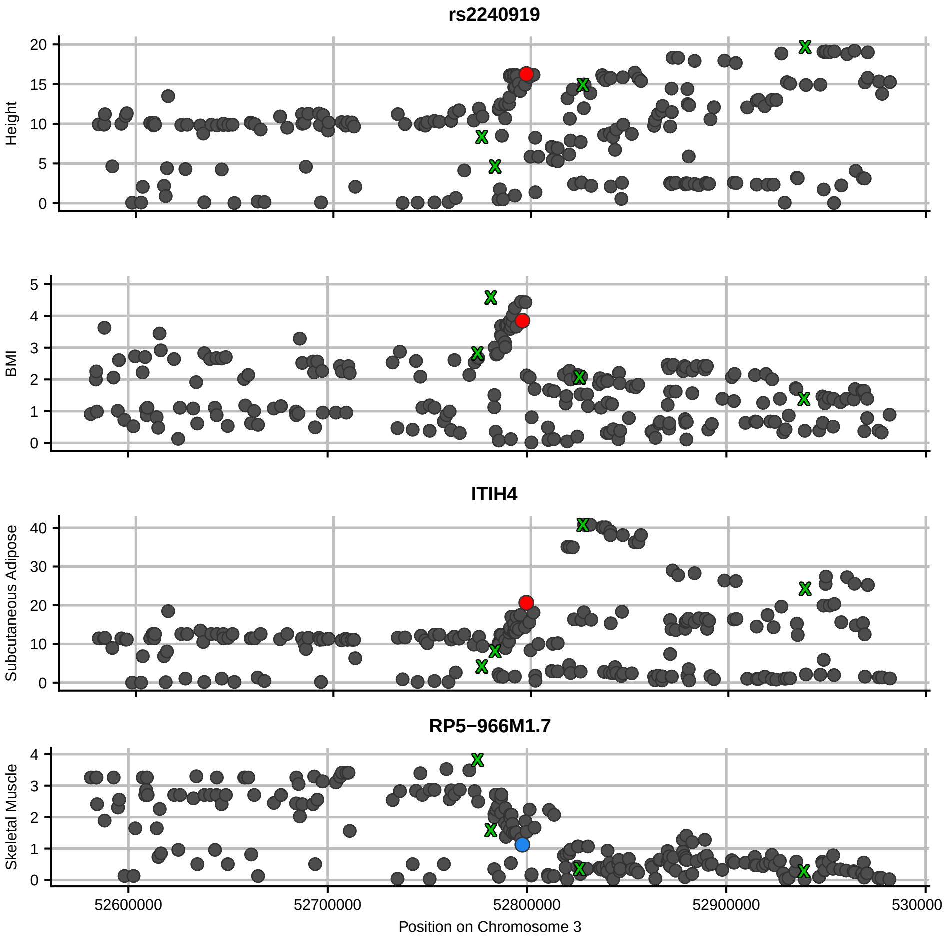

rs2240919
Height
BMI
ITIH4
Subcutaneous Adipose
RP5−966M1.7
Skeletal Muscle
52600000
52700000
52800000
52900000
530000
Position on Chromosome 3

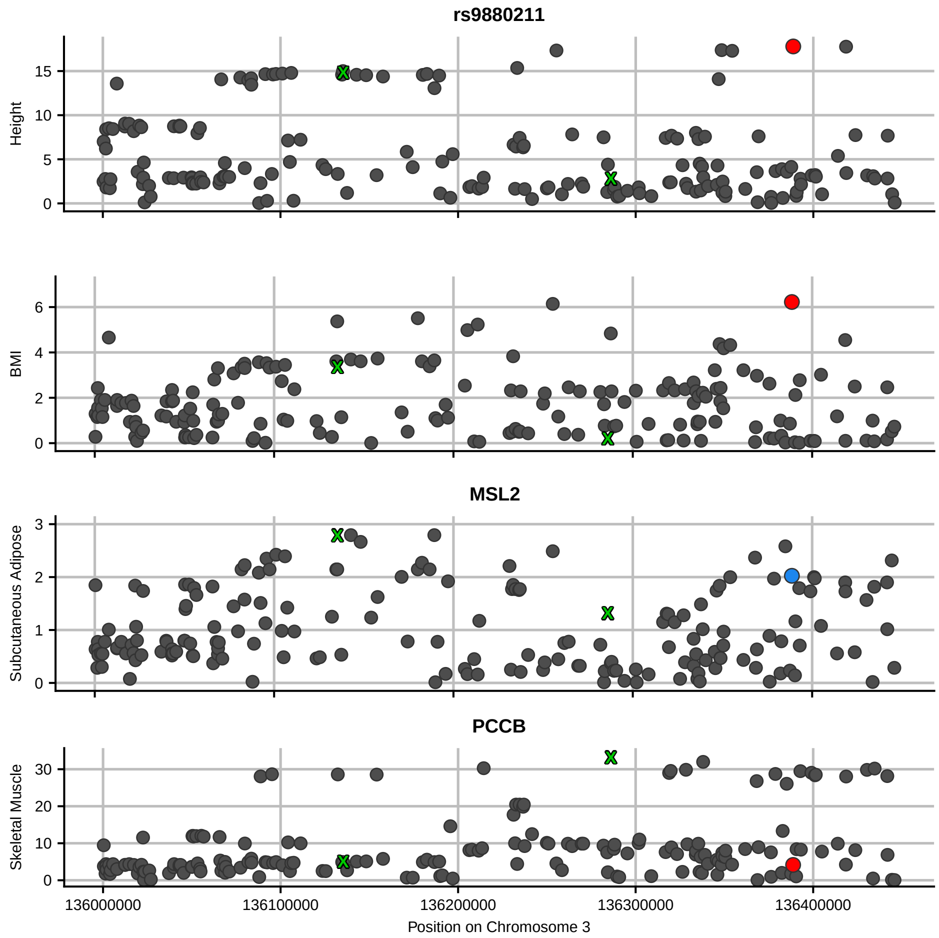

rs9880211
Height
BMI
MSL2
Subcutaneous Adipose
PCCB
Skeletal Muscle
136000000
136100000
136200000
136300000
136400000
Position on Chromosome 3

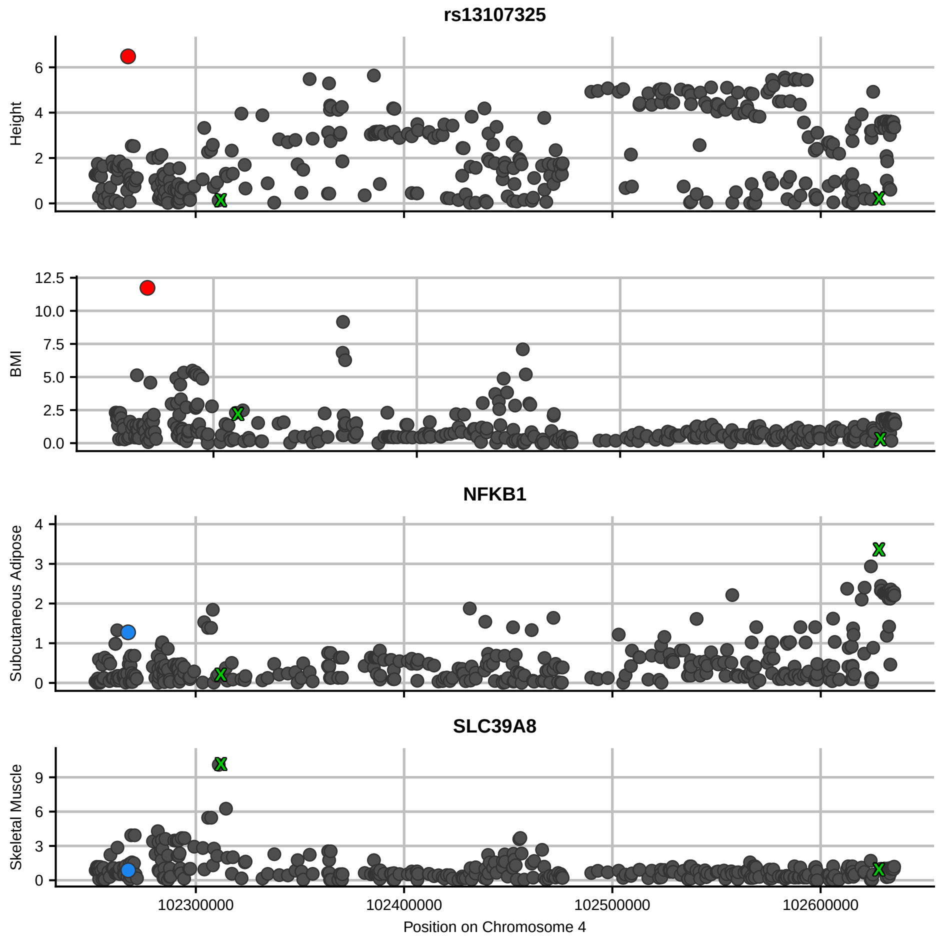
rs13107325
Height
BMI
NFKB1
Subcutaneous Adipose
SLC39A8
Skeletal Muscle
102300000
102400000
102500000
102600000
Position on Chromosome 4

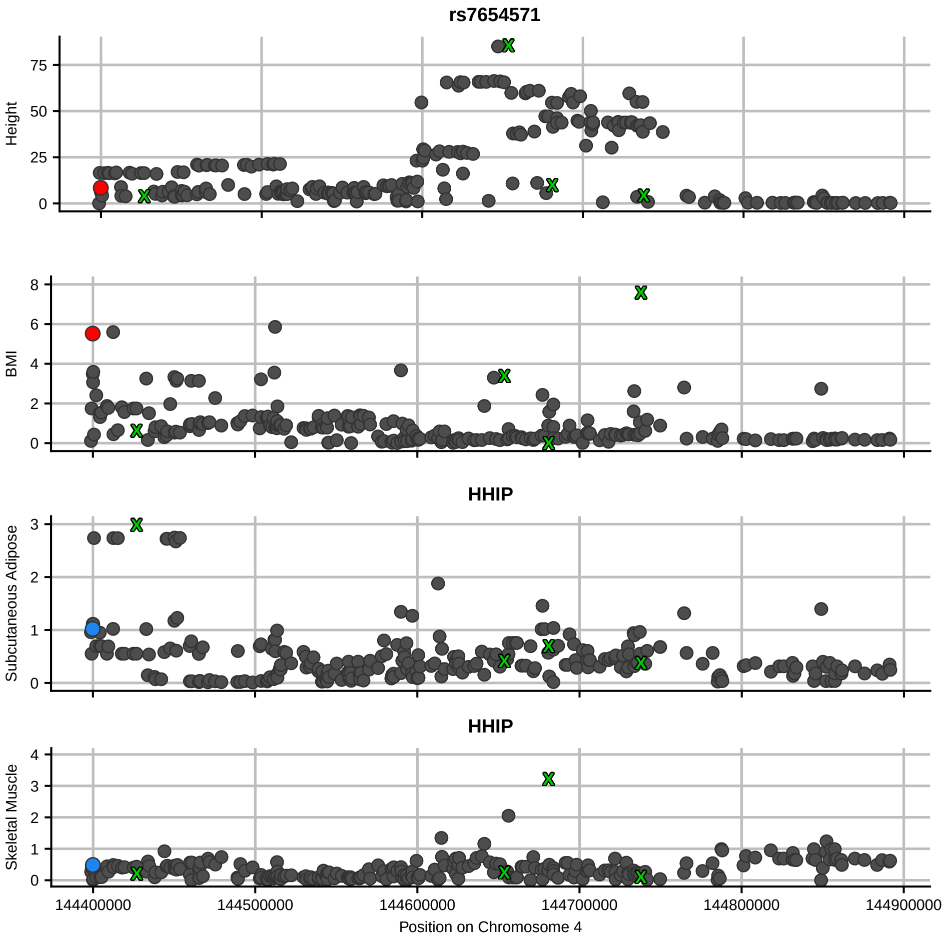

rs7654571
Height
BMI
HHIP
Subcutaneous Adipose
HHIP
Skeletal Muscle
144400000
144500000
144600000
144700000
144800000
144900000
Position on Chromosome 4

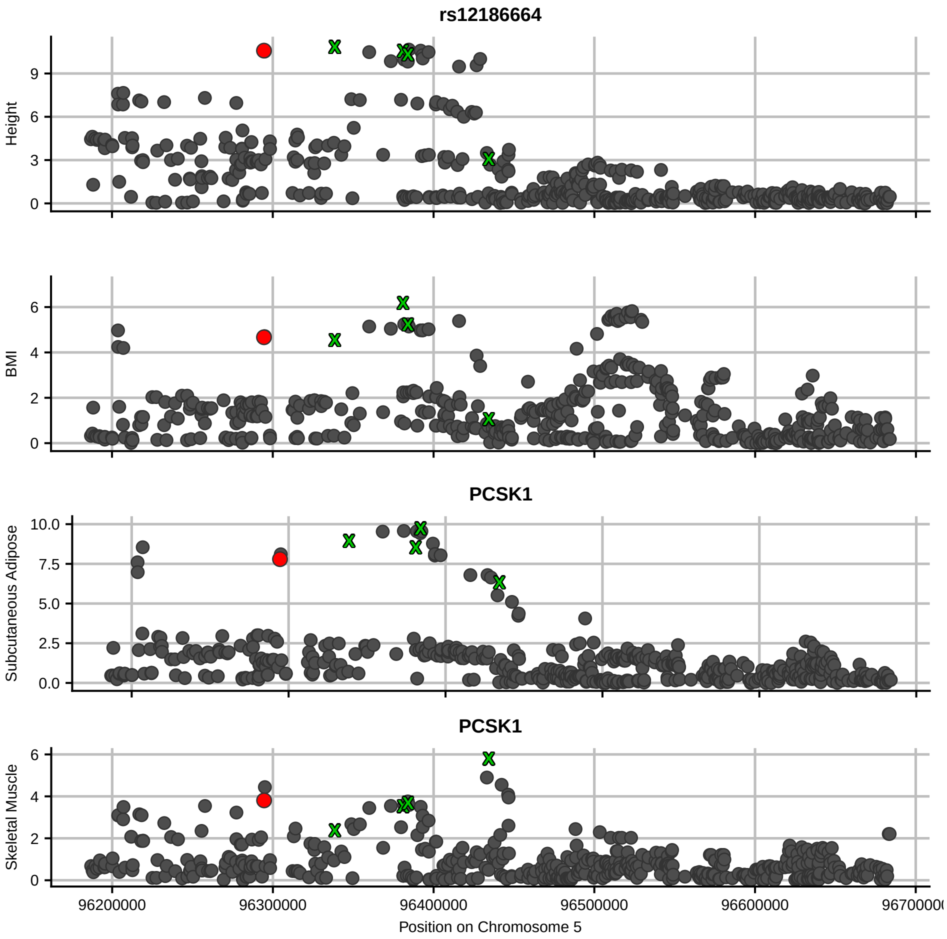
rs12186664
Height
BMI
PCSK1
Subcutaneous Adipose
PCSK1
Skeletal Muscle
96200000
96300000
96400000
96500000
96600000
Position on Chromosome 5

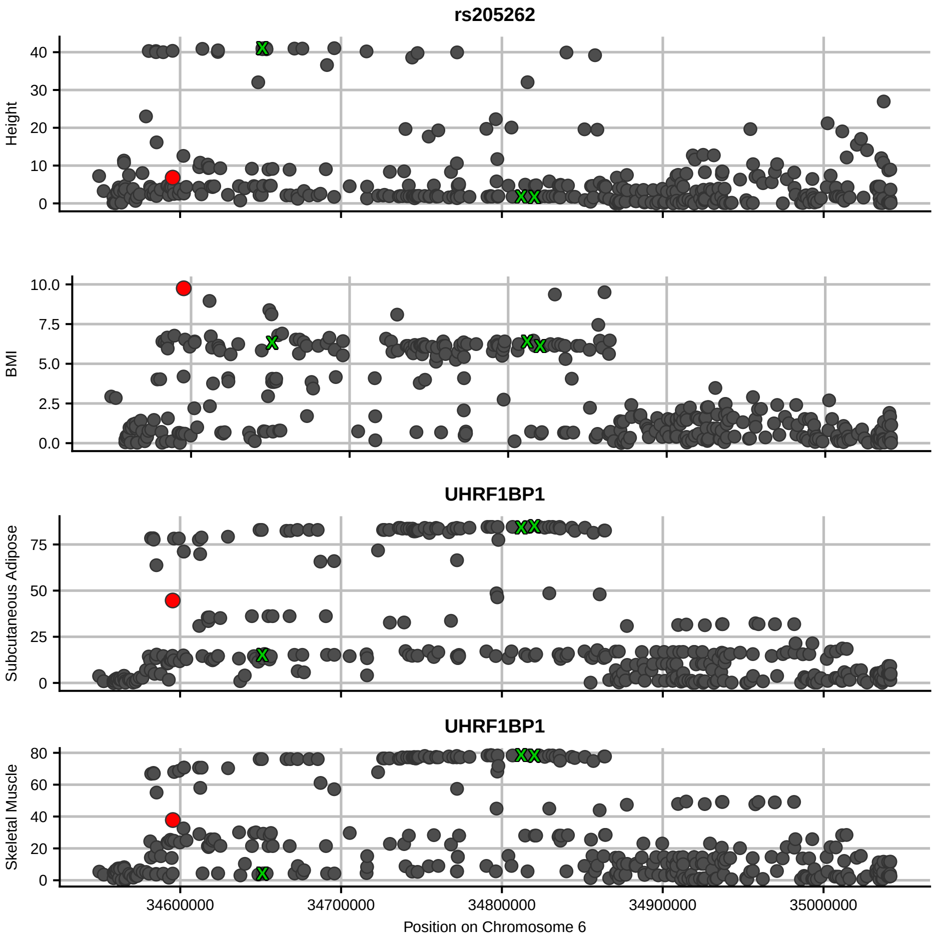

rs205262
Height
40
30
20
10
0
BMI
10.0
7.5
5.0
2.5
0.0
UHRF1BP1
Subcutaneous Adipose
75
50
25
0
UHRF1BP1
Skeletal Muscle
80
60
40
20
0
34600000
34700000
34800000
34900000
35000000
Position on Chromosome 6

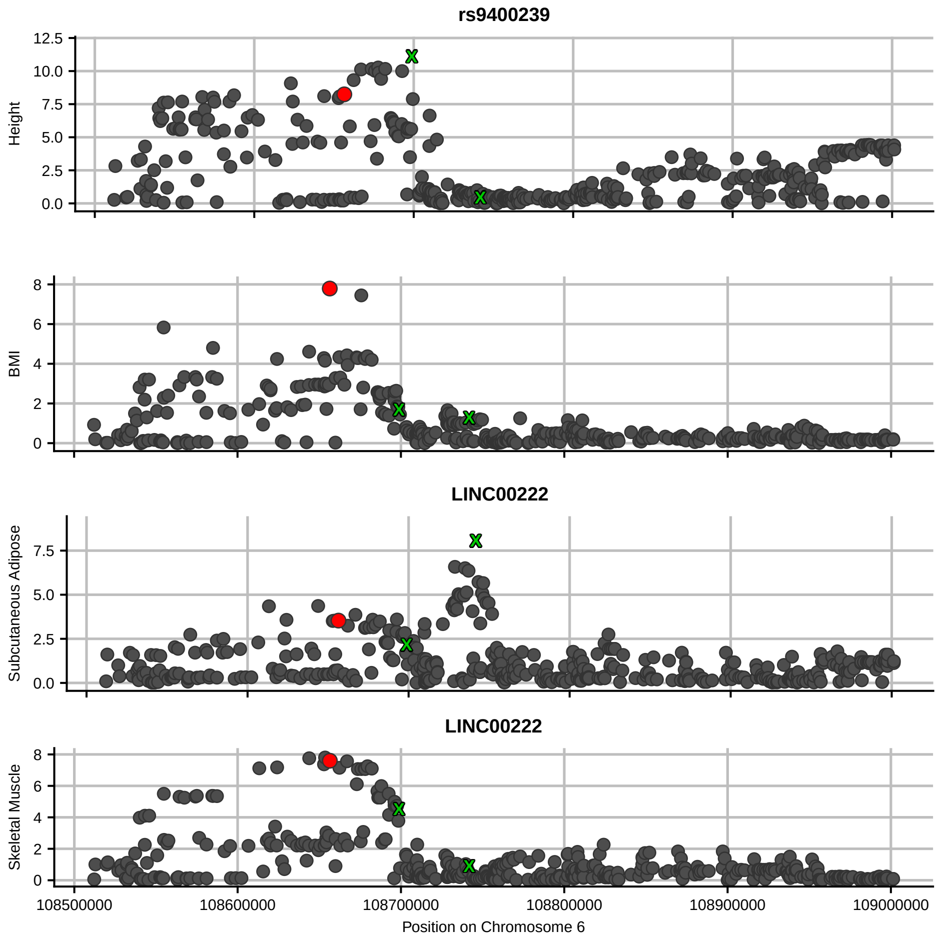

rs9400239
Height
BMI
LINC00222
Subcutaneous Adipose
LINC00222
Skeletal Muscle
108500000
108600000
108700000
108800000
108900000
109000000
Position on Chromosome 6

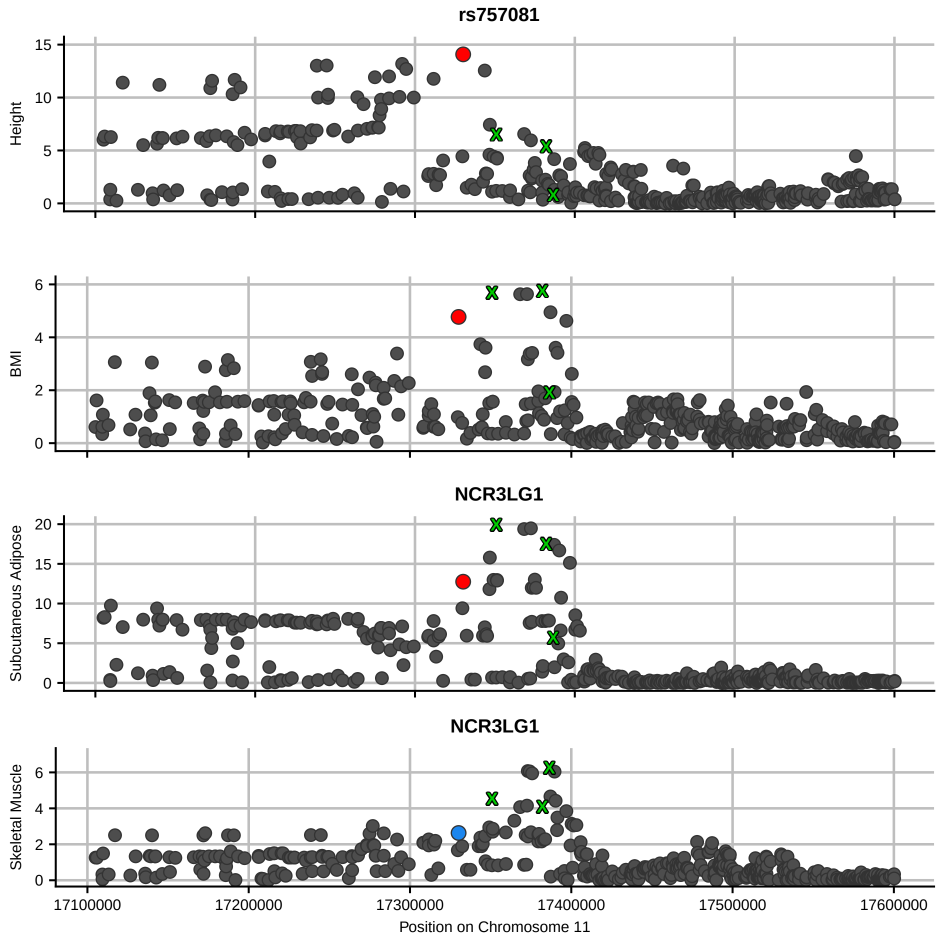

rs757081
Height
BMI
NCR3LG1
Subcutaneous Adipose
NCR3LG1
Skeletal Muscle
Position on Chromosome 11

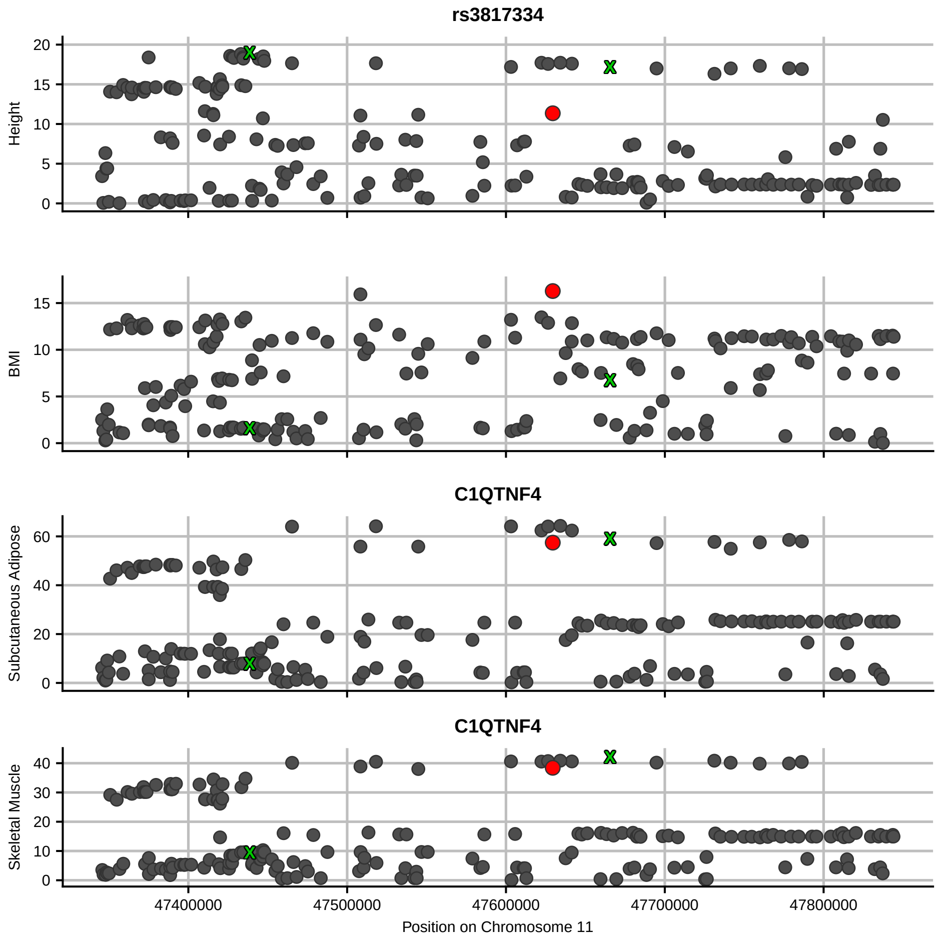

rs3817334
Height
20
15
10
5
0
BMI
15
10
5
0
C1QTNF4
Subcutaneous Adipose
60
40
20
0
C1QTNF4
Skeletal Muscle
40
30
20
10
0
47400000
47500000
47600000
47700000
47800000
Position on Chromosome 11

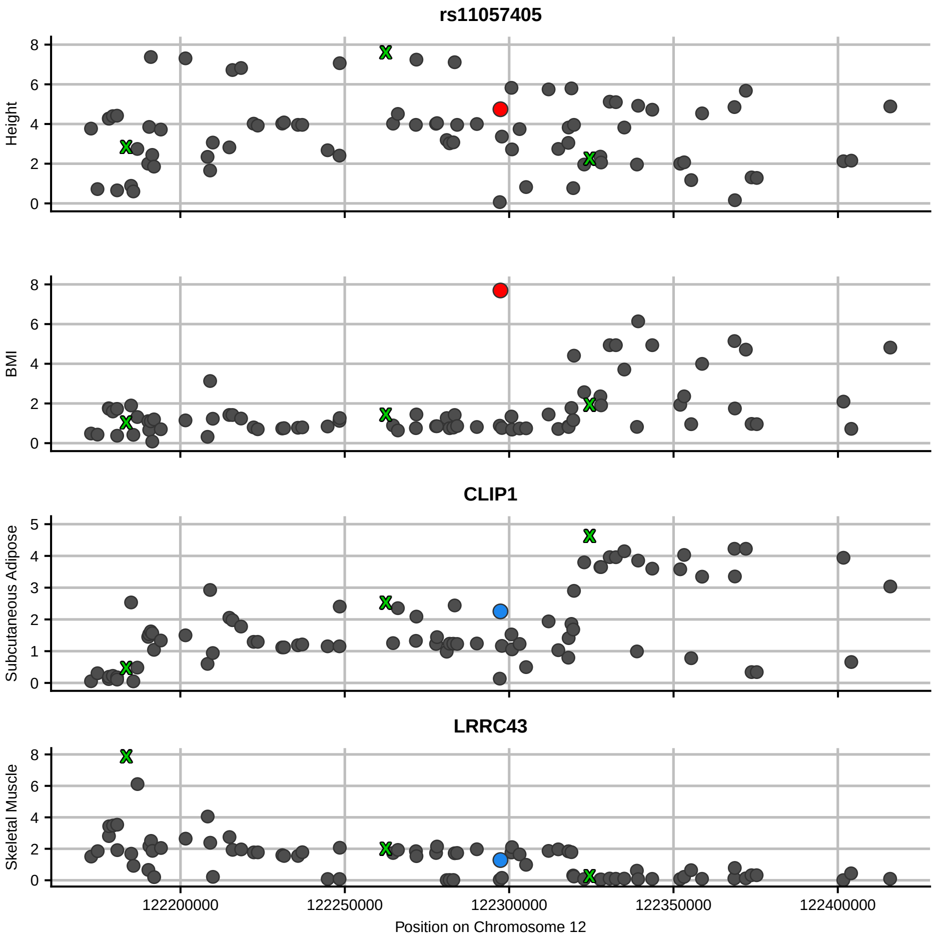

rs11057405
Height
BMI
CLIP1
Subcutaneous Adipose
LRRC43
Skeletal Muscle
122200000
122250000
122300000
122350000
122400000
Position on Chromosome 12

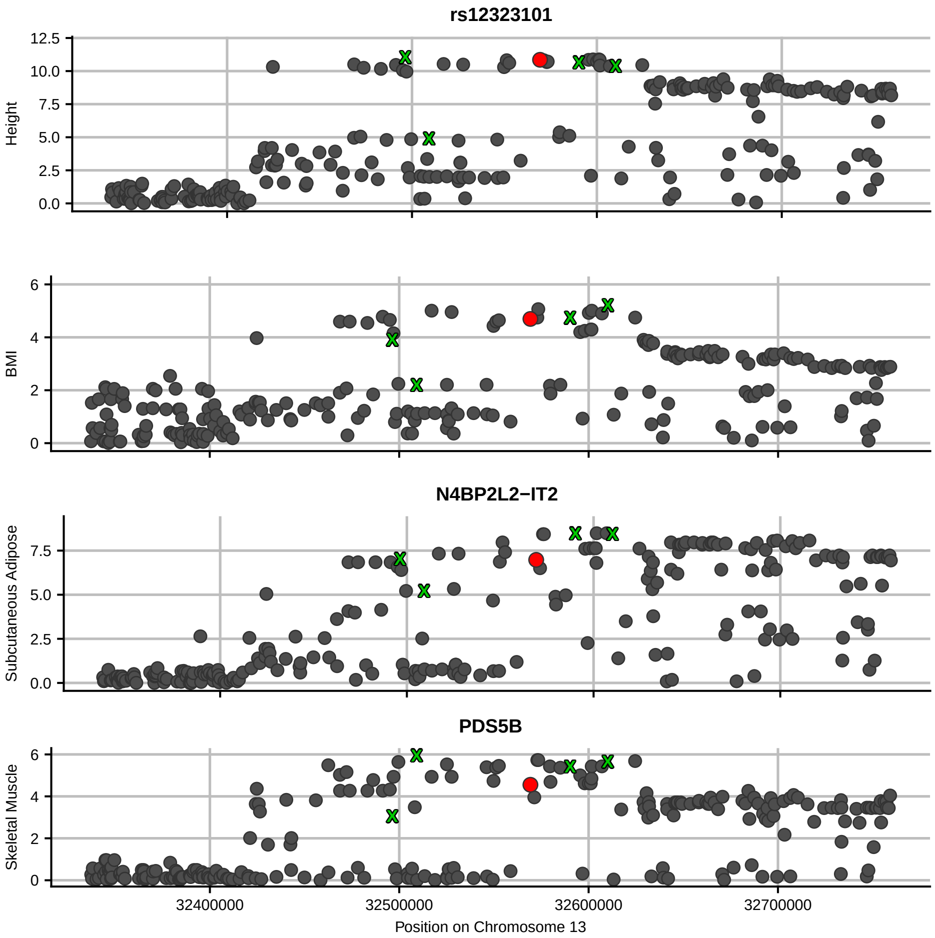
rs12323101
Height
BMI
N4BP2L2−IT2
Subcutaneous Adipose
PDS5B
Skeletal Muscle
32400000
32500000
32600000
32700000
Position on Chromosome 13

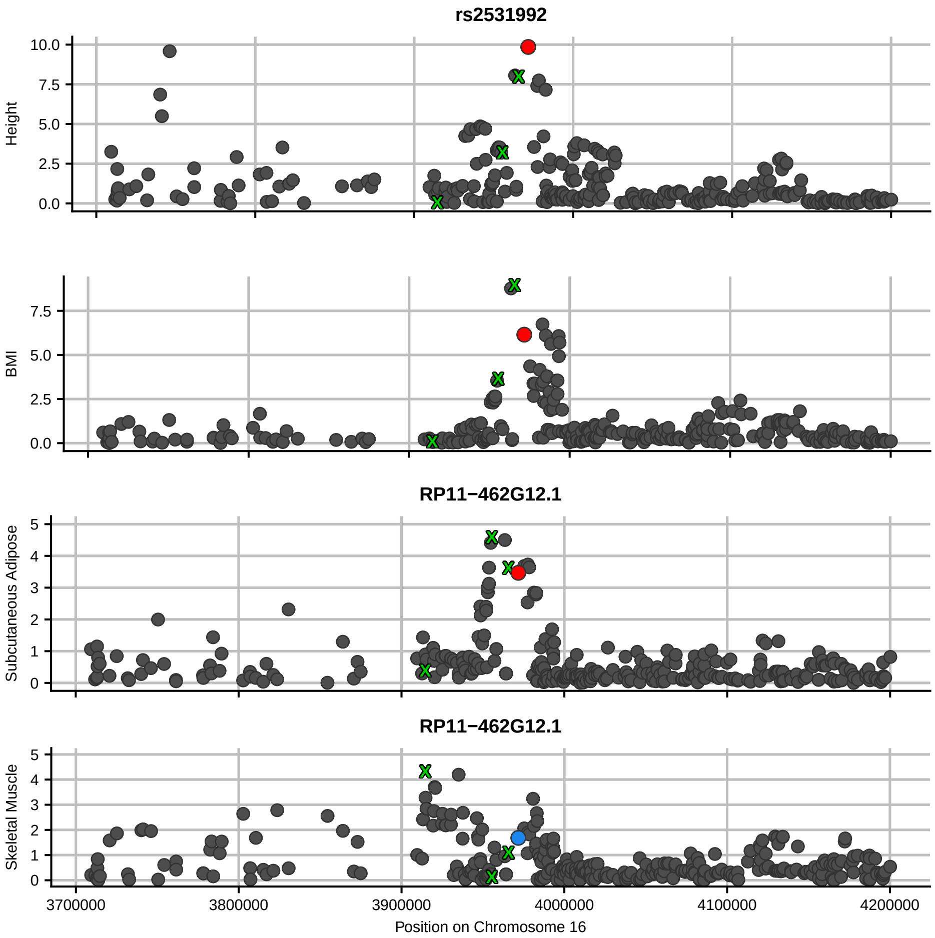

rs2531992
Height
BMI
RP11−462G12.1
Subcutaneous Adipose
RP11−462G12.1
Skeletal Muscle
Position on Chromosome 16
3700000
3800000
3900000
4000000
4100000
4200000

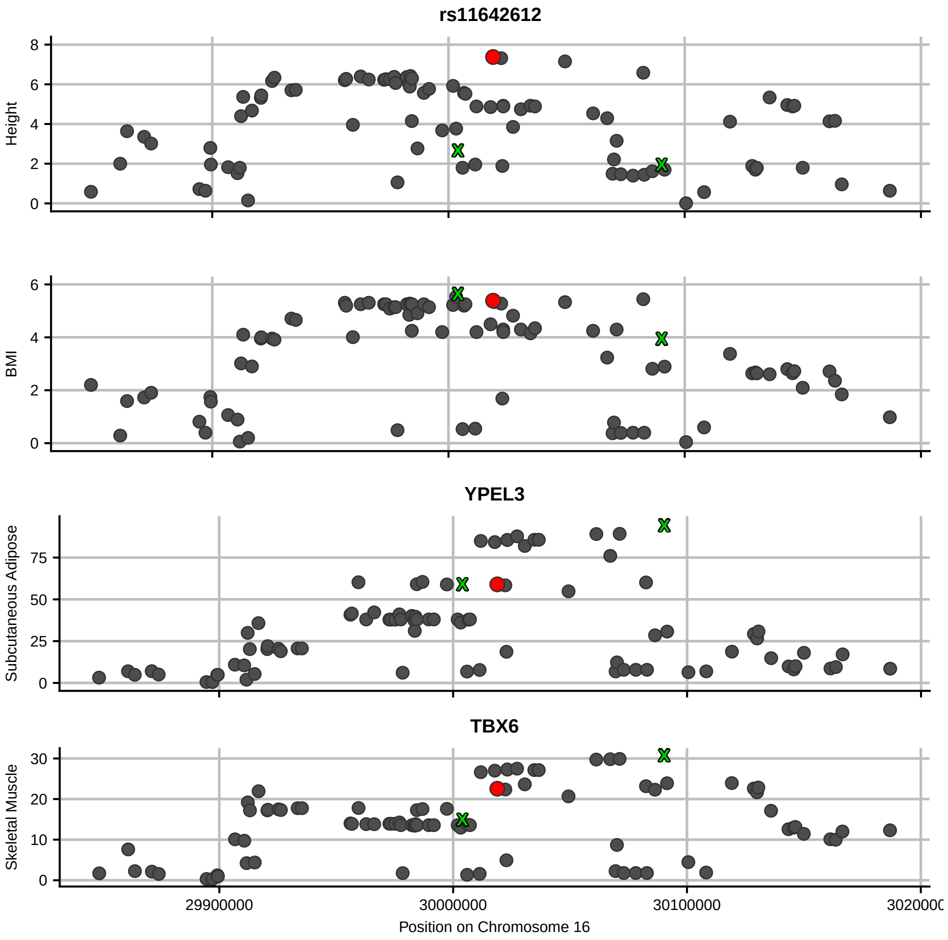

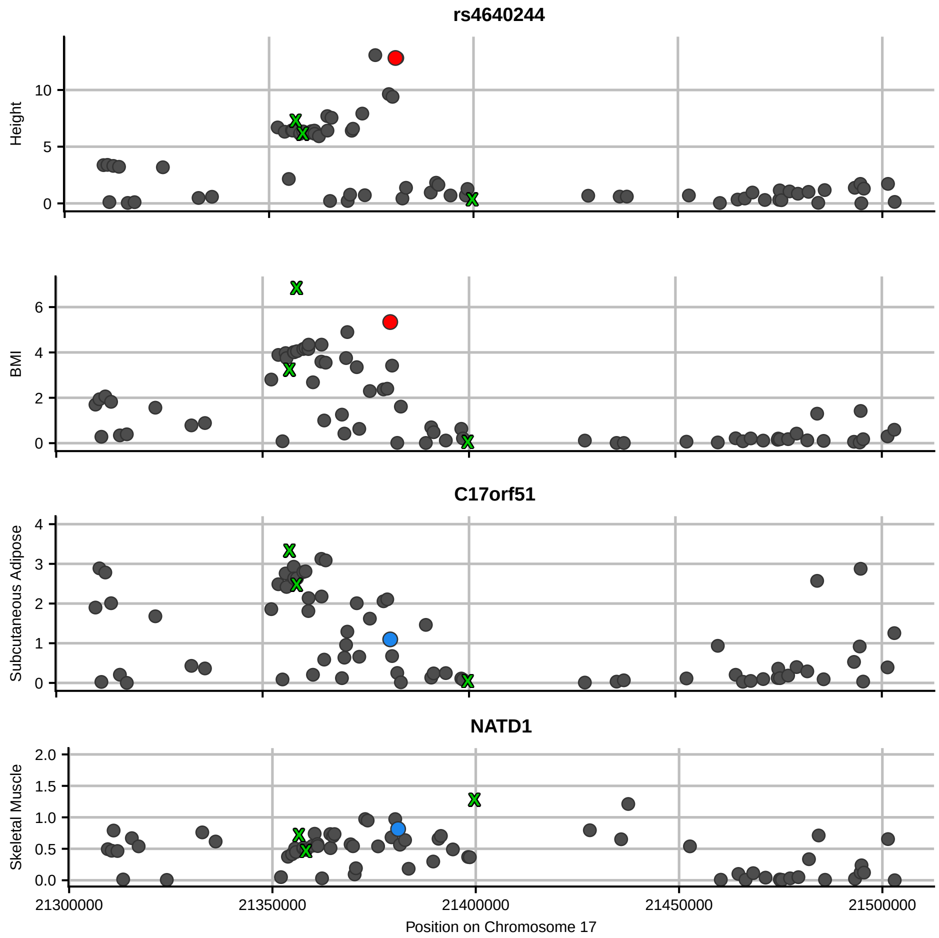
rs4640244
Height
BMI
C17orf51
Subcutaneous Adipose
NATD1
Skeletal Muscle
21300000
21350000
21400000
21450000
21500000
Position on Chromosome 17

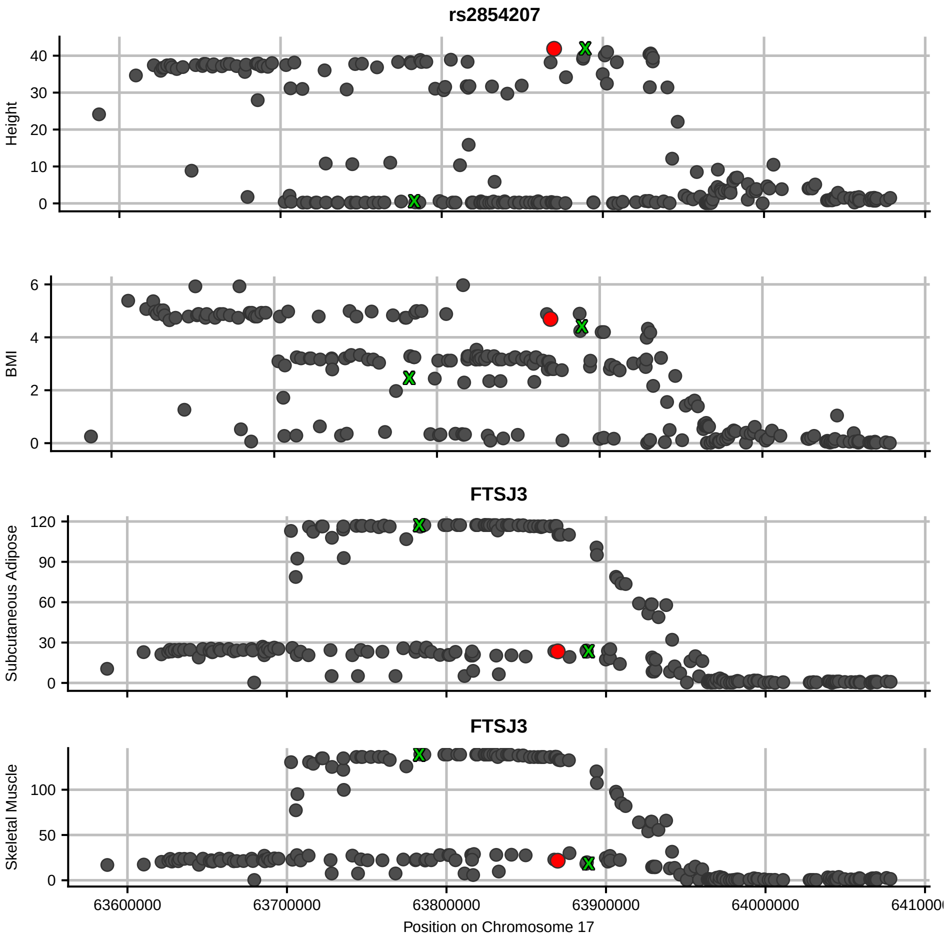
rs2854207
Height
BMI
FTSJ3
Subcutaneous Adipose
FTSJ3
Skeletal Muscle
63600000
63700000
63800000
63900000
64000000
Position on Chromosome 17

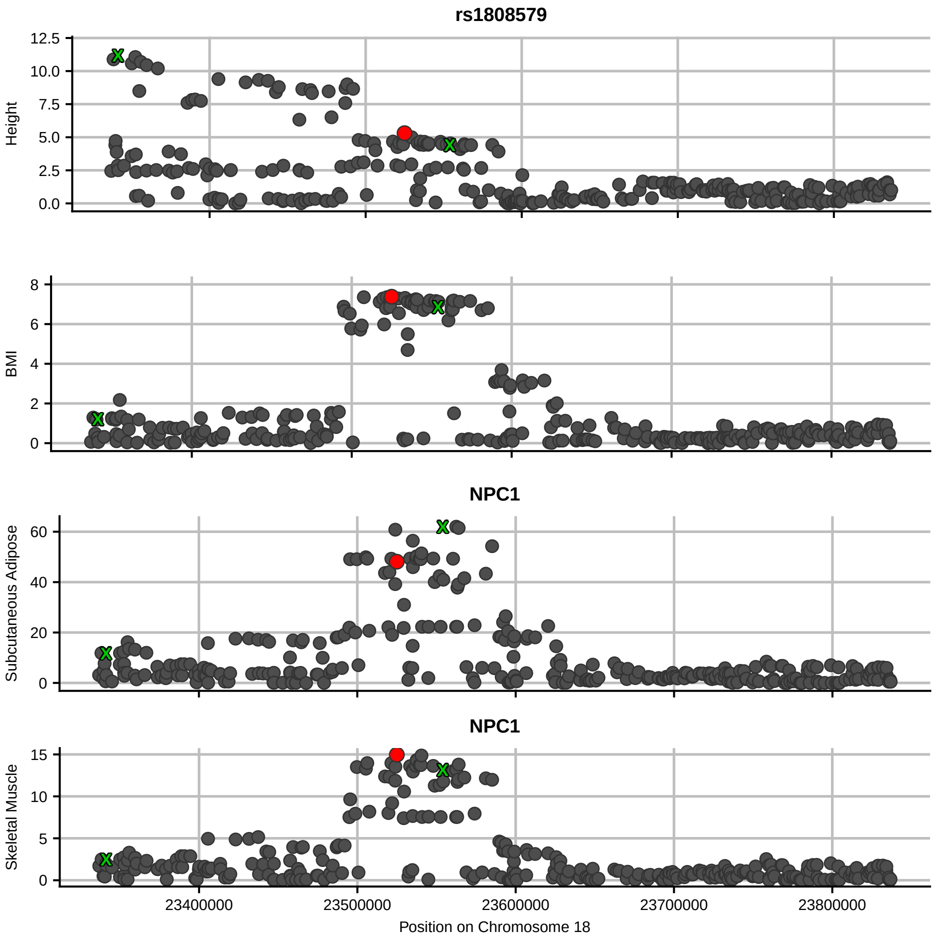

rs1808579
Height
12.5
10.0
7.5
5.0
2.5
0.0
BMI
8
6
4
2
0
NPC1
Subcutaneous Adipose
60
40
20
0
NPC1
Skeletal Muscle
15
10
5
0
23400000
23500000
23600000
23700000
23800000
Position on Chromosome 18

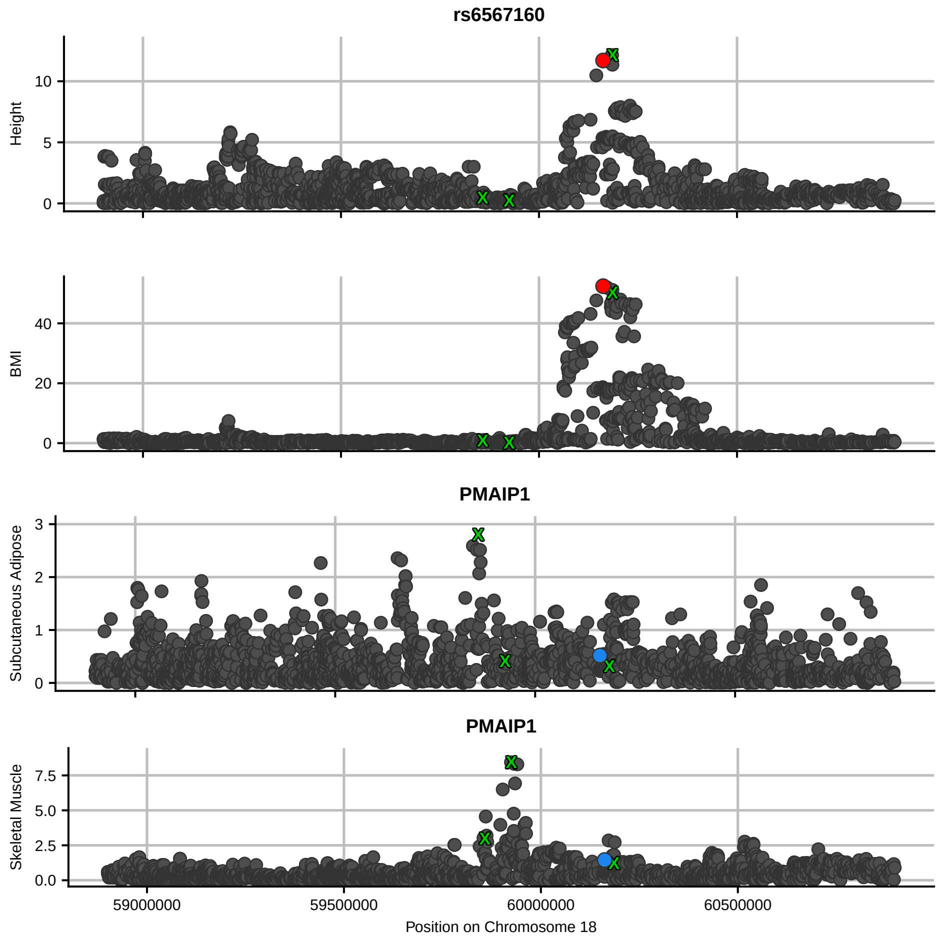
rs6567160
Height
BMI
PMAIP1
Subcutaneous Adipose
PMAIP1
Skeletal Muscle
59000000
59500000
60000000
60500000
Position on Chromosome 18

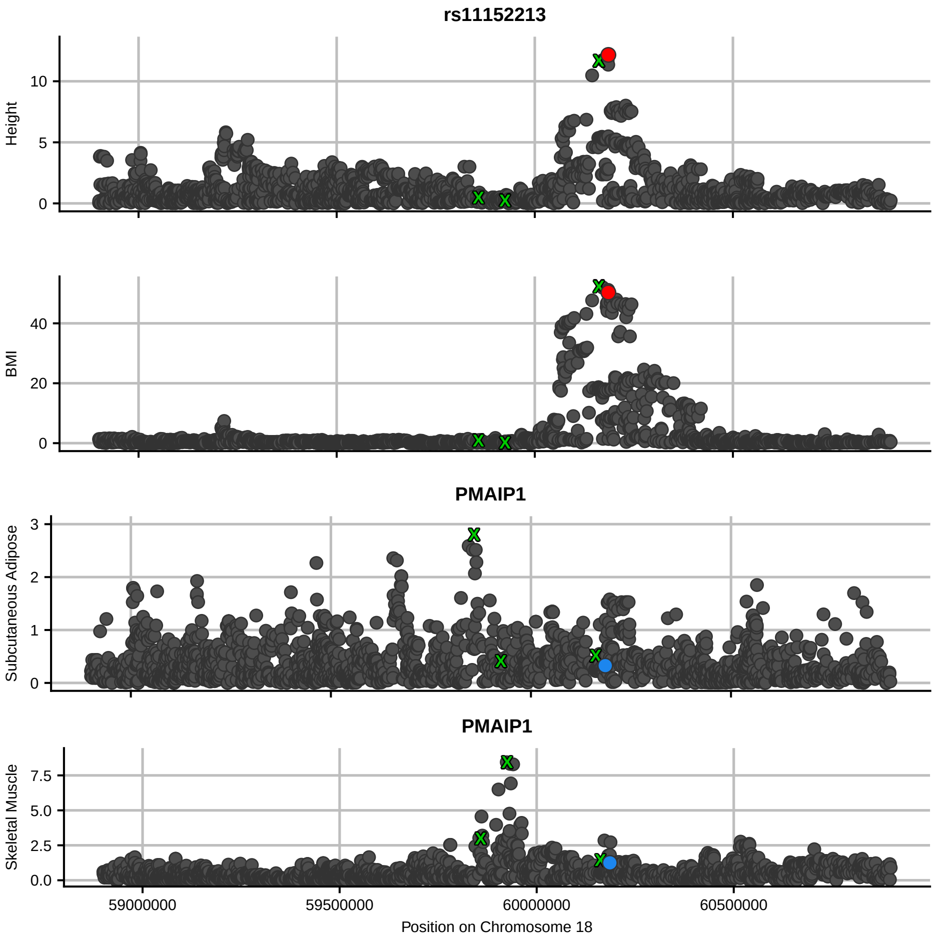

rs11152213
Height
BMI
PMAIP1
Subcutaneous Adipose
PMAIP1
Skeletal Muscle
59000000
59500000
60000000
60500000
Position on Chromosome 18

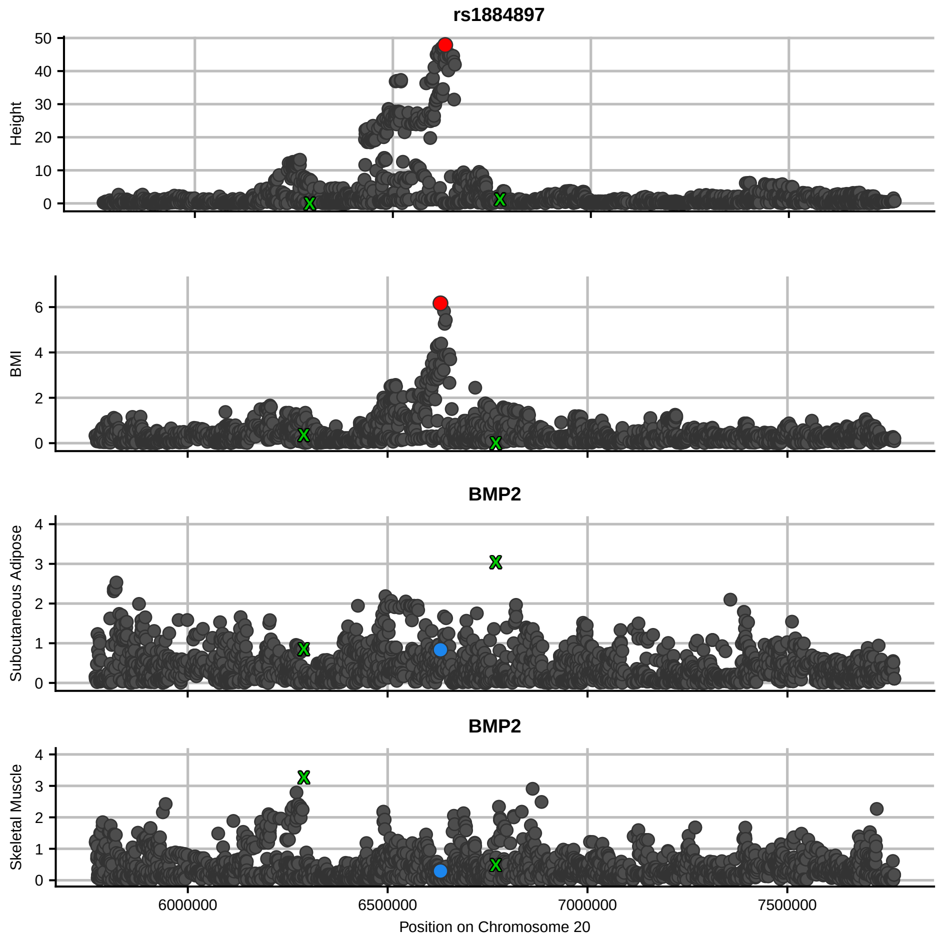

rs1884897
Height
BMI
BMP2
Subcutaneous Adipose
BMP2
Skeletal Muscle
6000000
6500000
7000000
7500000
Position on Chromosome 20
